# rsRNASP: A residue-separation-based statistical potential for RNA 3D structure evaluation

**DOI:** 10.1101/2021.09.20.461161

**Authors:** Ya-Lan Tan, Xunxun Wang, Ya-Zhou Shi, Wenbing Zhang, Zhi-Jie Tan

**Author notes:** Corresponding authors (to ZJT); (to WZ).

## Abstract

Knowledge-based statistical potentials have been shown to be rather effective in protein 3-dimensional (3D) structure evaluation and prediction. Recently, several statistical potentials have been developed for RNA 3D structure evaluation, while their performances are either still at low level for the test datasets from structure prediction models or dependent on the “black-box” process through neural networks. In this work, we have developed an all-atom distance-dependent statistical potential based on residue separation for RNA 3D structure evaluation, namely rsRNASP, which is composed of short- and long-ranged potentials distinguished by residue separation. The extensive examinations against available RNA test datasets show that, rsRNASP has apparently higher performance than the existing statistical potentials for the realistic test datasets with large RNAs from structure prediction models including the newly released RNA-Puzzles dataset, and is comparable to the existing top statistical potentials for the test datasets with small RNAs or near-native decoys. Additionally, rsRNASP is also superior to RNA3DCNN, a recently developed scoring function through 3D convolutional neural networks. rsRNASP and the relevant databases are available at website https://github.com/Tan-group/rsRNASP.

**SIGNIFICANCE:** RNAs play crucial roles in catalyzing biochemical reactions and regulating gene expression, and the biological functions of RNAs are generally coupled to their structures. Complementary to experiments, developing computational models to predict RNA 3D structures can be very helpful for understanding RNA biology functions. For a computational model, a reliable energy function is essentially important either for guiding conformational folding or for structure evaluation. For this purpose, we developed a residue-separation-based distance-dependent statistical potential, named rsRNASP which distinguishes the short- and long-ranged interactions, for RNA 3D structure evaluation. Our rsRNASP were examined against extensive test sets and shows overall superior performance over existing top traditional statistical potentials and a recently developed scoring function through 3D convolutional neural networks, especially for realistic test set from various computational structure prediction models.

## INTRODUCTION

Non-coding RNAs have critical biological functions in cell life activities, such as gene regulations and catalysis (1, 2). Generally, the functions of non-coding RNAs are coupled to their structures and consequently, understanding RNA structures, especially 3-dimensional (3D) structures, is crucial for understanding their biological functions (3, 4). However, due to the high cost and technical difficulty in experiment measurements, RNA 3D structures with high resolution deposited in protein data bank (PDB) database are still limited up to now (5). To complement experimental methods, various computational models have been developed to predict RNA 3D structures in silico (6, 7), including fragment-assembly-based models such as RNAComposer (8), MC-Fold/MC-Sym pipeline (9), Vfold3D (10, 11), 3dRNA (12, 13), and FARFAR (14, 15), and physics-based models such as iFold (16, 17), NAST (18), SimRNA (19), and a coarse-grained model with salt effect (20-25). Generally, a computational model for 3D structure prediction requires to involve an energy function, either for guiding conformational folding (26, 27), or for structure refinement (28) or for structure evaluation. Therefore, a reliable energy function is very important for the computational models for RNA 3D structure prediction.

Existing energy functions for protein and RNA 3D structure evaluation can be roughly grouped into two categories: physics-based force fields and knowledge-based statistical potentials (9, 12, 16, 19, 20, 29-43). Although physics-based force fields generally involve more lucid energy terms associated with bond lengths, bond angles, torsional angles, van der Waals and electrostatic interactions, they can become very inefficient or ineffective for large proteins or RNAs due to the required involvements of explicit solvent and salt. In contrast, knowledge-based statistical potentials derived from the experimental structures deposited in the PDB database have been shown to be very efficient and effective for the quality assessment of protein 3D structures, protein-protein and protein-ligand docking (44-54). The core difference between various statistical potentials mainly originates from the choice of different reference states (55), and up to now, six representative reference states have been proposed to build distance-dependent pairwise statistical potentials for proteins, i.e., averaging (48), quasi-chemical approximation (56), atom-shuffled (57), finite-ideal-gas (58), spherical-non-interacting (59) and random-walk-chain reference states (60).

For RNA 3D structure evaluation, several statistical potentials have been built based on different reference states. Bernauer et al have derived fully differentiable statistical potentials (KB) at both all-atom and coarse-grained levels, based on the quasi-chemical approximation reference state (61). Capriotti et al have built all-atom and coarse-grained statistical potentials (RASP) based on the averaging reference state (62). Wang et al have derived a combined distance- and torsion angle-dependent statistical potential (3dRNAscore) based on the averaging reference state, and 3dRNAscore emphasizes the local interactions through explicitly involving the backbone torsional angles (63). Zhang et al have proposed a distance-dependent statistical potential using finite-ideal-gas reference state (DFIRE-RNA) (64). In our recent work, for RNA 3D structure evaluation, we have made a comprehensive survey on the six existing reference states widely used for proteins with the same non-redundant RNA training set and parameters for RNA 3D structure evaluation (65). We have found that, the finite-ideal-gas and random-walk-chain reference states are modestly better than other ones in identifying native structures and ranking decoy structures and the existing traditional pair-wise statistical potentials only achieve a poor performance for realistic test dataset — RNA-Puzzles dataset (65). Beyond traditional pair-wise statistical potentials, Masso has proposed an all-atom four-body statistical potential to involve four-body effect in identifying native RNA structures (66). Recently, machine/deep learning approaches have also been applied to address this kind of problem (67, 68). Despite of “black-box” training/learning process, RNA3DCNN, a scoring function built by 3D deep convolutional neural networks, exhibits remarkably improved performance in identifying native structures for the realistic RNA-Puzzles test dataset whereas does not work so well for other test datasets composed of small RNAs or near-native decoys, compared with the existing top traditional statistical potentials (67). A newly released scoring function ARES from deep neural network based on training data from FARNAR showed good performance for evaluating structures from FARNAR while may has structure evaluation bias for FARNAR. Therefore, there is still lack of a reliable distance-dependent statistical potential with high performance for various structure prediction models.

Until now, most statistical potentials for RNAs do not distinguish the contributions of short-, medium- and long-ranged interactions. But for proteins, the explicit consideration of local (short- and medium-ranged) and nonlocal (long-ranged) interactions has been shown to be very helpful for understanding protein folding mechanism and predicting protein 3D structures (44, 69-75). Since RNA structure formation is generally hierarchical (76), the different residue-separation-ranged interactions may play different roles in RNA folding process or in stabilizing folded RNA structures. In RASP for RNAs, local interactions were separated from non-local ones with the residue (nucleotide) separation treated as topological factor, while the consideration of every residue separation for local interactions is too subtle to avoid sparse data problem, and RASP cannot achieve a good performance for RNA 3D structure evaluation (62).

In this work, we have developed an all-atom pair-wise distance-dependent statistical potential based on residue separation for RNA 3D structure evaluation, named rsRNASP, which is composed of short- and long-ranged potentials distinguished by residue separation. The extensive examinations show that rsRNASP has significantly improved performance than the existing statistical potentials for the realistic test datasets with large RNAs from various structure prediction models. It is encouraging that our traditional distance-dependent rsRNASP is overall superior to RNA3DCNN, a recently released scoring function through 3D convolutional neural networks.

## METHODS

### Residue separation and distance-dependent statistical potential

Since RNA structure folding is generally hierarchical (76), the different residue-separation-ranged interactions can play different roles in stabilizing folded RNA structures, and the potential energy of atomic pair interactions may strongly depend on their residue separations (45). Here, the residue separation *k* is defined by *k* = |*m* − *n*|, where *m* and *n* correspond to the observed residue sequence positions of a pair of atoms along an RNA chain. Thus, according to the inversed Boltzmann law, a pairwise distance-dependent statistical potential based on residue separation *k* can be given by (45)

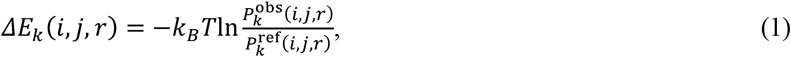

where *k*_*B*_ is the Boltzmann constant, and *T* is the Kelvin temperature. 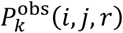 and 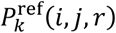 is the probability of the distance between atom pairs of types *i* and *j* with residue separation *k* in distance interval (*r, r* + *dr*] for native and reference states, respectively. In addition, the observed probability from native structure database can be obtained by

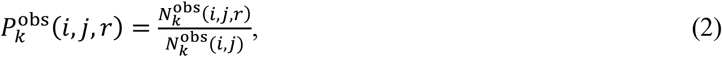

where 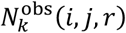 is the number of the distance between atom pair of atom types *i* and *j* with residue separation *k* in distance interval (*r, r* + *dr*], and 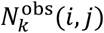 is the summation of 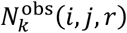 over *r*. As for 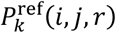, its form depends on the choice of simulated reference state, and as mentioned above, there have been six major reference states either based on the PDB database or based on various physical models (65).

### rsRNASP statistical potential based on residue separation

As discussed above, the classification of interaction ranges in a statistical potential may involve more accurate information extracted from native structure database (76). In developing rsRNASP, due to the severe sparse data problem arising from the current limited native RNA database, a residue separation threshold *k*_0_ was involved to distinguish between short- and long-ranged interactions instead of explicitly involving interactions at each residue-separation level. Through the statistical analyses on loop length distribution, *k*_0_ was taken as 4; see Section S2 and Fig. S1 in the Supporting Material for details. In addition, the local interactions between atoms within two residue intervals were ignored since such strong local interaction associated with local-chain connectivity restraints can overwhelm the other short- and long-ranged interactions essentially for stabilizing RNA global structures. Therefore, in rsRNASP, the interactions within 2 < *k* ≤ *k*_0_ were considered as the short-ranged interactions, and the energy for a conformation *C* of a given sequence *S* can be calculated by rsRNASP:

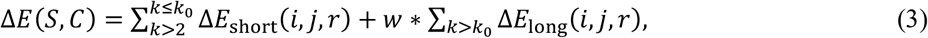

where

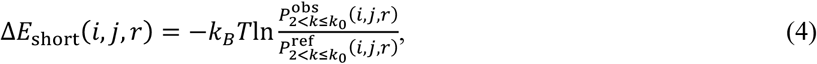

and

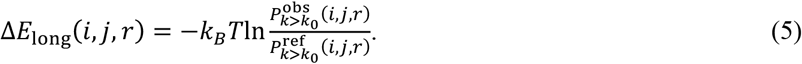

Here, *w* is a weight parameter to balance the contributions of short- and long-ranged interactions.

As shown in Eq. (1), the choice of reference states is crucial for building statistical potentials. Based on our previous work (65), in rsRNASP, we employed the random-walk-chain and averaging reference states to build long-ranged and short-ranged potentials since the random-walk-chain reference state has relatively good performance in identifying native structures and ranking decoy structures and the database-based averaging reference state may be complementary to those physical model-based ones (65). It is also noted that the random-walk-chain reference state with chain connectivity can directly distinguish interactions at different residue separations. The flow chart of building and testing steps of rsRNASP has been illustrated in Fig. 1.

**Figure 1.**
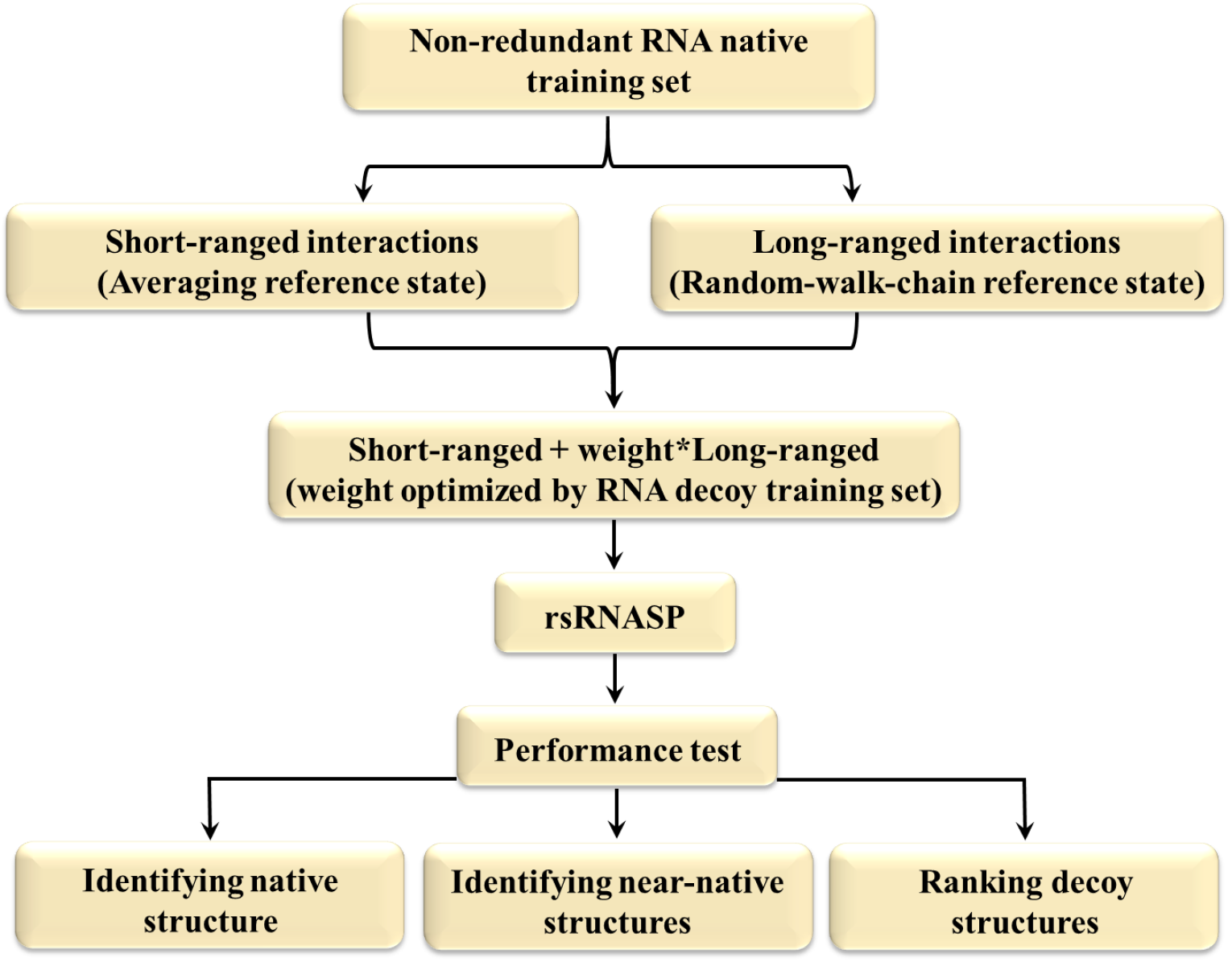
Flow chart of building and testing steps of rsRNASP.

### Training set and parameters

Beyond the training set in our previous work (65), we updated our non-redundant training set based on the RNA 3D Hub non-redundant set (77) (Release 3.102, 2019-11-27), which can be found in http://rna.bgsu.edu/rna3dhub/nrlist. First, 1641 representative RNA chains with an X-ray resolution < 3.5 Å were collected from RNA 3D Hub list (Release 3.102). Afterwards, we removed the chains whose structures are complexed with proteins or DNAs and those with chain length ≤10 nt. Finally, using BLASTN program (78), we removed the RNA structures with sequence identity > 80% and coverage > 80%. Through the prior operation steps, our training set contains 191 RNA structures and the PDB IDs of these 191 RNAs were presented in Table S1 in the Supporting Material. It should be noted that a small amount of RNAs in the test sets have sequence identity > 80% and coverage > 80% with the native RNAs in the training set, and these RNAs were still kept in the training set for maintaining the full structure spectrum (65). For these RNAs, the leave-one-out or jackknife method was used for testing the performance of rsRNASP (62, 65).

Moreover, according to previous works (63, 65) and Section S3 as well as Figs. S2 and S3 in the Supporting Material, the distance bin width was taken as 0.3 Å, and the distance cutoffs were set to 13 Å and 24 Å for short- and long-ranged potentials in rsRNASP, respectively. For the situation that some atom pairs were not observed within a certain bin width, the potentials were set to the highest potential value in the whole potential range of corresponding atom pair types, and *k*_*B*_*T* was taken as the unit of potential energy in this work.

To derive the weight *ω* between short- and long-ranged interactions, we used a training decoy set to optimize it. First, 35 single-stranded RNAs in our training set were selected, including wide structure spectrum of hairpin loops, internal loops, junction loops and tertiary interactions; see the PDB IDs of these 35 RNAs in Table S1 in the Supporting Material. Afterwards, for the 35 RNAs, we employed four RNA 3D structure prediction models (FARFAR2 (15), RNAComposer (8), SimRNA (19) and 3dRNA v2.0 (13)) to generate about 10 decoy structures for each RNA by giving secondary structures parsed by x3DNA-dssr (79) from the webservers with default output (FARFAR2 (15), RNAComposer (8) and 3dRNA v2.0 (13)) or first frames of clusters (obtained with a 3.5Å RMSD threshold) of a simulated conformation trajectory (SimRNA (19)), thus about 40 decoy structures were generated for each RNA. Simultaneously, an RNA length *N*-dependent function *f*(*N*) was involved to normalize the *N*-dependent atom-pair number of the long-ranged interactions due to the large residue-separation range and the consequent *N*-dependent atom-pair number, and 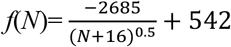 based on the statistical analyses on the training native set. Consequently, *w* = *w*_0_/*f*(*N*) and *w*_0_ was chosen to be 16, according to the extensive examinations on the training decoy dataset. Please see Section S4 and Figs. S4 and S5 in the Supporting Material for the details of weight *w*.

### Test datasets

To evaluate the performance of rsRNASP and make comparisons with other existing statistical potentials, similar to previous works (63, 67), three test sets were used in this work, including a new test subset built by ourselves.

Test set I, called randstr decoy set (62), consists of 85 RNAs with decoy structures generated by MODELLER (80) with a set of Gaussian restraints for atom distances and dihedral angles, and there are 500 decoy structures for each native RNA. Test set I can be downloaded from http://melolab.org/supmat/RNApot/Sup._Data.html. Test set II is composed of decoys built by Bernauer et al (61) and Das et al (14). The former includes two subsets (61): MD subset consists of 5 RNAs with about 3500 decoy structures for each RNA generated by replica-exchange molecular dynamics simulations with atom position restrained, and NM subset consists of 15 RNAs with 500 decoy structures for each RNA generated by normal mode perturbation method. The third subset called FARNA subset is composed of 20 RNAs with about 500 decoy structures for each RNA, which was generated by FARNA (14).

Test set III is composed of RNA-Puzzles_standardized (81) and prediction-model subsets (PM subset). The former subset was obtained from RNA-Puzzles, which is a CASP-like evaluation of blind 3D RNA structure predictions (81). RNA-Puzzles_standardized subset contains 22 different puzzles and dozens of decoy structures for each native RNA from different RNA structure prediction models and can serve as a realistic test set for illustrating the performance of a statistical potential in evaluating RNA 3D structures. The older version of RNA-Puzzles (Puzzles_normalized) including only 18 puzzles, may have smaller number of decoys for some puzzles compared with the updated version. Since the existing RNA statistical potentials were almost often assessed against the older version, the test on normalized-submissions was also presented in this work. Furthermore, for more extensive examination against realistic decoy structures, we built a new test subset, named prediction-models (PM) subset. Specifically, to ensure the sequence identify < 80% and coverage < 80% with RNAs in our training set, 20 single-stranded RNAs were filtered from recently updated PDB database (5) (2021-03-20). In accordance with the way of assembling the training decoy set, given the sequences and secondary structures, 10 decoy structures were generated for each RNA by four RNA 3D structure models (FARFAR2 (15), RNAComposer (8), SimRNA (19) and 3dRNA v2.0 (13)). Thus, PM subset consists of 20 RNAs with about 40 decoy structures for each native RNA. The RNA-Puzzles_standardized dataset can be downloaded from https://github.com/RNA-Puzzles/standardized_dataset and the PM subset can be found in https://github.com/Tan-group/rsRNASP.

### Measuring RNA structure similarity

To measure the structural similarity between two RNA structures, the metrics of DI (deformation index) was used in addition to root-mean-square-deviation (RMSD), and DI is defined as (83):

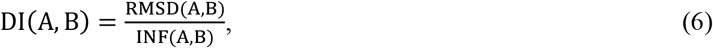

where RMSD(A, B) and INF(A, B) reflects the difference of geometry and topology between structures A and B, respectively. INF(A, B) is the interaction network fidelity between two structures A and B, and is measured by Matthews correlation coefficient of base-pairing and base-stacking interactions between A and B (84). If two structures occupy the most same interactions, the value of DI would be similar to RMSD, and otherwise, DI value would be relatively larger than RMSD. In this work, these two metrics of DI and RMSD were both used, and the tools for calculating DI and INF can be downloaded from https://github.com/RNA-Puzzles/BasicAssessMetrics (85).

## RESULTS AND DISCUSSION

### Evaluation metrics

Generally, the ability of identifying native and near-native structures from corresponding decoy ones and the ability of ranking the decoy structures reasonably can describe the performance of a statistical potential (63). Therefore, we used the number of native structures with the lowest energy, the number of native structures within top five ones of the lowest energies, enrichment score (ES) (61, 63-65, 67) and Pearson correlation coefficient (PCC) (62, 65) calculated by a statistical potential for test sets as evaluation metrics. ES is defined as (61):

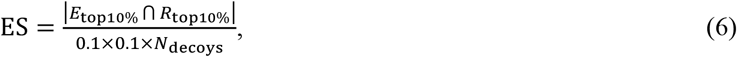

where |*E*_top10%_ ⋂ *R*_top10%_| is the intersection between the top 10% near-native structures (measured by DI or RMSD) and the top 10% of the structures with the lowest energies. *N*_decoys_ is the total number of decoy structures for one native RNA. The value of ES ranges from 1 to 10, and 10 represents the best performance. PCC is given by (62):

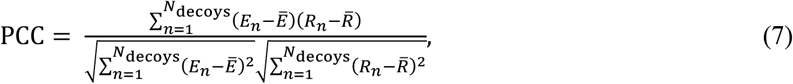

where *E*_*n*_ and *R*_*n*_ are the energy and DI (or RMSD) of the *n*th structure, respectively. Ē and 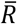 are the average energy and DI (or RMSD) of decoys, respectively. The value of PCC ranges from 0 to 1. If the relationship between energies and DIs (or RMSDs) is completely linear (PCC is equal to 1), the statistical potential would have a perfect performance.

In addition, since there are not enough predicted structures (decoys) for each RNA in RNA-Puzzles_standardized and PM subsets, the DI (or RMSD) of structure with the lowest energy, and the rank of the nearest-native structure (DI or RMSD) were used instead of ES values.

### Performance of rsRNASP on test set I

Test set I consists of 85 RNAs with decoy structures generated by MODELLER (62), and the RMSDs of decoy structures are mainly distributed in a narrow range of [0-6 Å] (65), suggesting that the decoy structures from the perturbation method are almost near-native ones. As shown in Table 1, rsRNASP can identify 82 native structures out of 85 RNA decoy sets. Thus, in identifying native structures, rsRNASP is apparently superior to RNA3DCNN and is slightly better than RASP while is slightly weaker than DFIRE-RNA and 3dRNAscore. RNA3DCNN, RASP, 3dRNAscore and DFIRE-RNA identify 63, 80, 84 and 85 native structures out of decoy ones of 85 RNAs, respectively. In fact, the three native structures unidentified by rsRNASP generally have very low energies close to the lowest ones. For a further evaluation, we made statistics on the number of native structures identified within top five ones of the lowest energies for the statistical potentials. As also shown in Table 1, rsRNASP could identify 85 out of 85, i.e., rsRNASP, DFIRE-RNA and 3dRNAscore have the same top performance. In contrast, RASP and RNA3DCNN identify 84 and 79 out of 85, respectively.

**Table 1.**
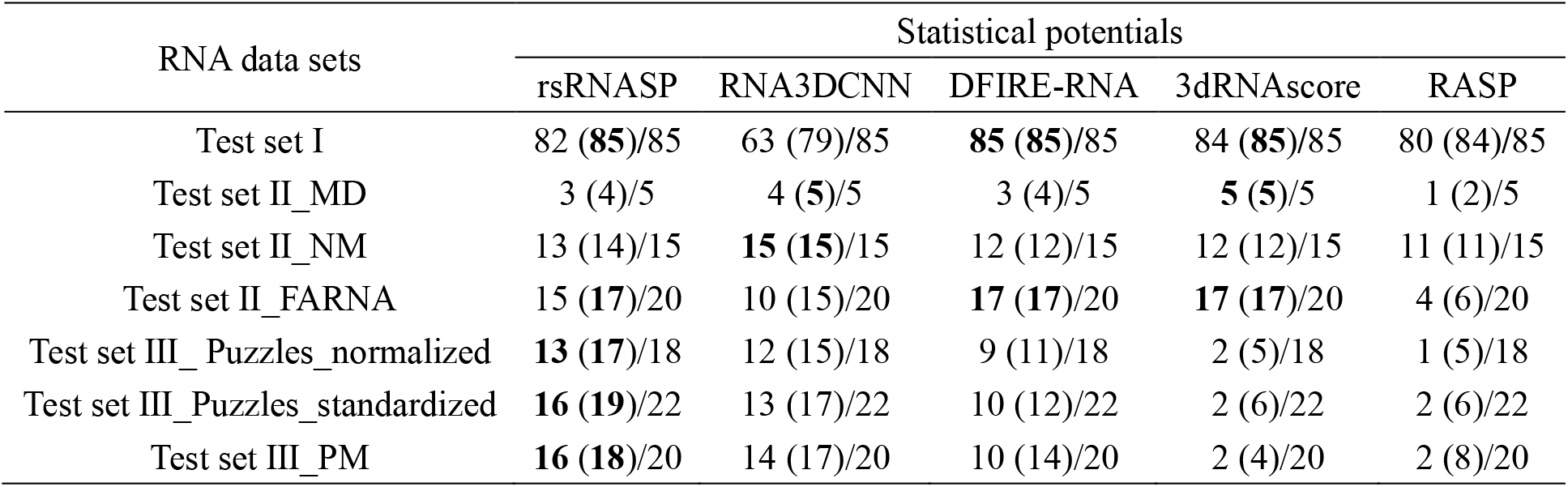
The number of identified native structures and the number of structures identified within top five ones of the lowest energies (in brackets) by rsRNASP and other existing statistical potentials.

Furthermore, we calculated the average ES and PCC values for test set I. As shown in Table S2 in the Supporting Material, the average ES values on DIs obtained by rsRNASP, RNA3DCNN, DFIRE-RNA, 3dRNAscore and RASP are 8.6, 8.9, 8.7, 9.0 and 9.0, respectively, and the average PCC values on DIs are 0.80, 0.84, 0.81, 0.82, 0.86, respectively. The ES and PCC values indicate that for test set I, the correlations between DIs and energies are all very strong for those five statistical potentials which all reach high performance. Additionally, we calculated average ES and PCC values on RMSDs show the similar trend to those on DIs; see Table S2 in the Supporting Material. The RMSD-energy scatterplots for all the 85 RNAs in test set I by rsRNASP were shown in Fig. S6 in the Supporting Material.

Therefore, for test set I, the performance of our rsRNASP is very close to the top ones of the existing statistical potentials (e.g., DFIRE-RNA and 3dRNAscore) and is also at high level in identifying native and near-native structures and in ranking decoy structures.

### Performance of rsRNASP on test set II

MD subset composed of 5 RNAs was produced by replica-exchange molecular dynamics simulations (61) and the RMSDs of the decoy structures are distributed in a very narrow range of [0-3 Å] (65), suggesting that the decoy structures are very close to their native ones. For MD subset, as shown in Table 1, 3 native structures from the decoy ones of 5 RNAs are identified by rsRNASP, which is similar to DFIRE-RNA and superior to RASP while is slightly weaker than 3dRNAscore and RNA3DCNN. Furthermore, we also calculated the number of native structures identified within top five ones of the lowest energies for the subset. As shown in Table 1, rsRNASP identifies 4 out of 5 decoy sets, and its performance is superior to RASP while is still slightly lower than RNA3DCNN and 3dRNAscore. Actually, the native RNA unidentified by rsRNASP is 1nuj, and its energy is ranked to the sixth in the corresponding decoy set. On the other hand, we calculated the ES and PCC values for MD subset. As shown in Table S3 in the Supporting Material, the average ES value on DIs calculated by rsRNASP is ∼7.1, which is higher than that of RASP (∼6.4) and similar to that of DFIRE-RNA (∼7.1) and RNA3DCNN (∼7.2) while is slightly lower than that of 3dRNAscore (∼7.9). Furthermore, the average PCC values on DIs obtained by rsRNASP, RNA3DCNN, DFIRE-RNA, 3dRNAscore and RASP are 0.81, 0.73, 0.82, 0.75 and 0.74, respectively. Additionally, the ES and PCC values calculated on RMSDs show the similar trend to those on DIs; see Table S4 in the Supporting Material. Therefore, for MD subset in test set II, the performance of our rsRNASP is very close to the top ones of the existing statistical potentials (e.g., 3dRNAscore for identifying near-native structures and DFIRE-RNA in ranking decoy structures). The RMSD-energy scatter-plots of all five decoy sets in MD subset were provided in Fig. S7 in the Supporting Material.

NM subset consisting of decoy structures for 15 RNAs was generated by normal-mode perturbation method (61) and the decoy structures are very close to the native ones, as reflected by the very narrow RMSD range of [1-4 Å] (65). For NM subset, as shown in Table 1, rsRNASP identifies 13 native structures out of 15 decoy sets, and is slightly superior to the existing traditional statistical potentials including DFIRE-RNA, 3dRNAscore and RASP while is slightly weaker than RNA3DCNN. It is noted that these two native structures unidentified by our rsRNASP are 1esy and 1kka, which were solved by NMR spectroscopy at low salt. The low salt solutions would cause the less compact structures for RNAs due to the polyanionic nature of RNAs (86-88). As shown in Table S3 in the Supporting Material, the average ES values on DIs calculated by rsRNASP, RNA3DCNN, DFIRE-RNA, 3dRNAscore and RASP are 5.6, 6.1, 5.4, 6.4 and 5.3, respectively, and the average PCC values for five statistical potentials are 0.89, 0.88, 0.91, 0.90 and 0.87, suggesting the high performance of these five statistical potentials for NM subset. Additionally, the ES and PCC values on RMSDs show the very similar trend to those on DIs; see Table S4 in the Supporting Material. Therefore, for NM subset, our rsRNASP is very close to the top statistical potentials (e.g., RNA3DCNN) in identifying native structures and ranking decoy structures (e.g., DFIRE-RNA). The RMSD-energy scatter-plots of all decoy sets in NM subset were provided in Fig. S7 in the Supporting Material.

FARNA subset composed of decoy structures of 20 RNAs, was generated by FARNA (14), and the RMSDs of decoys are dispersed in the wide range of [1-15 Å] (65). For FARNA subset, as shown in Table 1, rsRNASP, RNA3DCNN, DFIRE-RNA, 3dRNAscore and RASP identify 15, 10, 17, 17, and 4 native structures out of 20 RNA decoy sets, respectively. Thus, rsRNASP can identify apparently more native structures than RNA3DCNN and RASP, while appears slightly worse than DFIRE-RNA and 3dRNAscore. However, Table 1 shows that 17 native RNAs can be identified within top five ones of the lowest energies by rsRNASP, i.e., the two unidentified native structures in the 17 RNAs also have very relatively low energies calculated by rsRNASP. Thus, rsRNASP has the similar performance to DFIRE-RNA and 3dRNAscore and better performance than RNA3DCNN and RASP in identifying native structures. As shown in Table S3 in the Supporting Material, for ES and PCC values on DIs, though RNA3DCNN performs the best than other statistical potentials, these five statistical potentials all have unsatisfactory performances with overall low mean ES values and PCC values. This may be attributed to that the energy landscapes composed of the decoy conformations are very rugged with some local energy minimums in addition to the native structures with lowest energies. The ES and PCC values on RMSDs were also shown in Table S4 in the Supporting Material and the full RMSD-energy scatter-plots of all decoy sets for FARNA subset were provided in Fig. S7 in the Supporting Material.

### Performance of rsRNASP on test set III

Test set III is composed of Puzzles_standardized and PM subsets which are generated from various RNA 3D structure prediction models and contains large RNAs with large RMSD range, and consequently test set III can serve as a realistic test set for a statistical potential beyond test sets I and II composed of small RNAs or near-native decoy structures mainly from perturbation methods.

RNA-Puzzles_standardized subset consists of dozens of decoy structures from the different 3D structure prediction models for 22 different RNA puzzles and was generally considered as a realistic test set for a statistical potential in evaluating RNA 3D structures (64, 67, 81). For the subset, as shown in Table 1, our rsRNASP can identify 16 native structures out of decoys for 22 RNAs, while RNA3DCNN, DFIRE-RNA, 3dRNAscore and RASP identify 13, 10, 2 and 2, respectively. For native structures identified within top five ones of the lowest energies, rsRNASP, RNA3DCNN, DFIRE-RNA, 3dRNAscore and RASP can identify 19, 17, 12, 6 and 6 out of 22 decoy sets, respectively; see Table 1. Moreover, as shown in Table 2, the average DIs of structures with the lowest energies from rsRNASP, RNA3DCNN, DFIRE-RNA, 3dRNAscore and RASP are 4.6, 5.9, 7.6, 17.1 and 17.8, respectively. As for PCC values on DIs, rsRNASP has a higher average PCC value of 0.57, compared with those from DFIRE-RNA (∼0.52), RNA3DCNN (∼0.35), 3dRNAscore (∼0.35) and RASP (∼0.38). Therefore, overall, for RNA-Puzzles_standardized subset, our rsRNASP has the apparently better performance in identifying native structures and in ranking decoy structures, compared with the other statistical potentials including RNA3DCNN from neural network. The ES and PCC values on RMSDs were shown in Table S5 in the Supporting Material, and as an illustrative example, the relationships between RMSDs and energies calculated by these five scoring functions for puzzle-10 on which rsRNASP revealed the highest PCC value were respectively shown in the Fig. S8 in the Supporting Material. Moreover, the full RMSD-energy scatter-plots of all decoy sets in Puzzles_standardized subset obtained by rsRNASP were also given in Fig. S9 in the Supporting Material. In addition, for a historic comparison with the existing statistical potentials, we also show the evaluations against the Puzzles_normalized dataset in Table 1, as well as Table S6 and Table S7 in the Supporting Material, and the full RMSD-energy scatter-plots for Puzzles_normalized subset were given in Fig. S10 in the Supporting Material. The evaluations indicate that rsRNASP also has apparently better performance than the existing statistical potentials for Puzzles_normalized dataset.

**Table 2.**
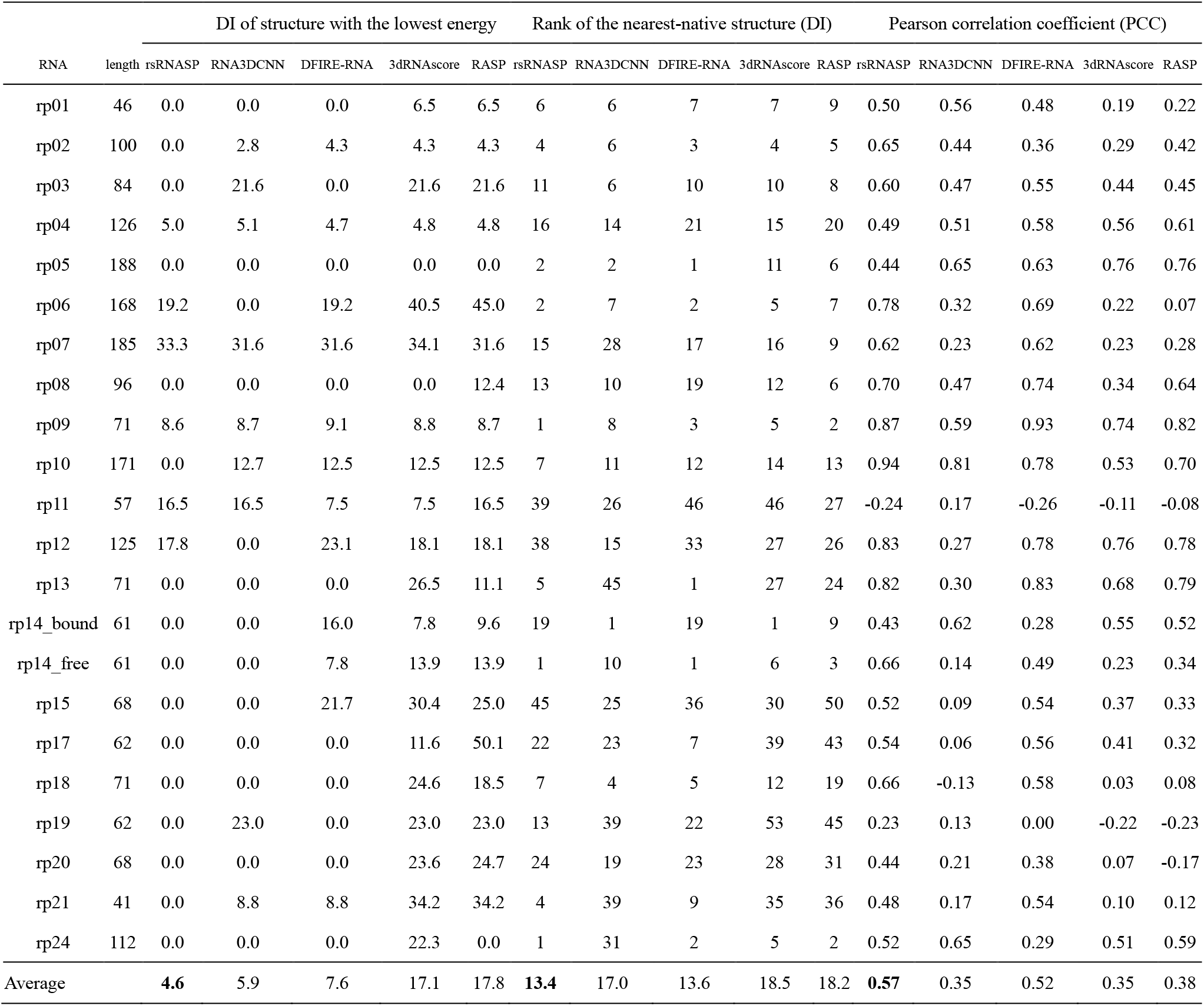
The DIs of structures with the lowest energy, ranks of the nearest-native structure (DI) and Pearson correlation coefficients between energies and DIs of decoy structures calculated by rsRNASP and other statistical potentials for Puzzles_standardized subset in test set III.

PM subset is composed of decoys from four RNA 3D structure prediction models for 20 single-stranded RNAs. For PM subset, as shown in Table 1, rsRNASP can identify 16 native structures out of 20 RNA decoy sets, while RNA3DCNN, DFIRE-RNA, 3dRNAscore and RASP identify 14, 10, 2 and 2, respectively. Furthermore, Table 1 also shows that, for native structures identified within top five ones of the lowest energies, rsRNASP can identify 18 RNAs, while RNA3DCNN, DFIRE-RNA, RASP and 3dRNAscore can identify 17, 14, 8 and 4 out of 20, respectively. Furthermore, as shown in Table 3, for this subset, rsRNASP gives the average DI of the structures with the lowest energy of 3.3 and the average PCC value of 0.64, which is apparently better than those from other statistical potentials. The evaluations based on RMSDs show the similar trend to those based on DIs; see Table S8 in the Supporting Material. As a typical example, the relationships between RMSDs and energies obtained by all five scoring functions for 3RKF on which rsRNASP has a high PCC value were respectively shown in the Fig. S8 in the Supporting Material. The full RMSD-energy scatter-plots of all decoy sets in PM subset calculated by rsRNASP were given in Fig. S11 in the Supporting Material.

**Table 3.** The DIs of structures with the lowest energy, ranks of the nearest-native structure and Pearson correlation coefficients between energies and DIs of decoy structures calculated by the different statistical potentials for test set III_PM subset.

| RNA | length | DI of structure with the lowest energy |  |  |  |  | Rank of the nearest-native structure |  |  |  |  | Pearson correlation coefficient (PCC) |  |  |  |  |
| --- | --- | --- | --- | --- | --- | --- | --- | --- | --- | --- | --- | --- | --- | --- | --- | --- |
|  |  | rsRNASP | RNA3DCNN | DFIRE-RNA | 3dRNAscore | RASP | rsRNASP | RNA3DCNN | DFIRE-RNA | 3dRNAscore | RASP | rsRNASP | RNA3DCNN | DFIRE-RNA | 3dRNAscore | RASP |
| 1Z43 | 101 | 0.0 | 27.4 | 32.7 | 23.6 | 15.1 | 22 | 33 | 27 | 18 | 24 | 0.66 | 0.53 | 0.49 | 0.03 | 0.07 |
| 3A3A | 86 | 4.8 | 4.2 | 15.5 | 30.0 | 13.9 | 6 | 5 | 14 | 14 | 18 | 0.85 | 0.90 | 0.63 | 0.65 | 0.64 |
| 3IVN | 69 | 0.0 | 0.0 | 0.0 | 8.3 | 0.0 | 1 | 5 | 6 | 26 | 28 | 0.48 | 0.49 | 0.31 | 0.14 | -0.16 |
| 3L0U | 73 | 0.0 | 0.0 | 0.0 | 15.8 | 10.7 | 1 | 2 | 16 | 20 | 26 | 0.80 | 0.76 | 0.76 | -0.34 | -0.53 |
| 3LA5 | 71 | 0.0 | 0.0 | 0.0 | 0.0 | 30.5 | 1 | 1 | 1 | 15 | 14 | 0.88 | 0.57 | 0.87 | 0.04 | -0.36 |
| 3RKF | 67 | 0.0 | 0.0 | 1.1 | 12.9 | 15.4 | 3 | 1 | 6 | 11 | 3 | 0.95 | 0.69 | 0.97 | 0.47 | 0.48 |
| 3SKI | 68 | 0.0 | 0.0 | 0.0 | 11.8 | 14.0 | 2 | 3 | 6 | 12 | 7 | 0.89 | 0.80 | 0.82 | 0.23 | 0.21 |
| 4AOB | 94 | 0.0 | 0.0 | 0.0 | 14.9 | 22.8 | 1 | 1 | 1 | 23 | 17 | 0.78 | 0.69 | 0.69 | -0.39 | -0.34 |
| 4FEN | 67 | 0.0 | 0.0 | 0.0 | 12.0 | 35.4 | 2 | 3 | 2 | 14 | 12 | 0.86 | 0.59 | 0.88 | 0.04 | -0.30 |
| 5D5L | 77 | 0.0 | 0.0 | 0.0 | 24.2 | 14.3 | 10 | 6 | 11 | 14 | 20 | 0.72 | 0.47 | 0.54 | 0.41 | 0.38 |
| 5FJC | 93 | 0.0 | 0.0 | 0.0 | 25.8 | 0.0 | 1 | 7 | 1 | 19 | 6 | 0.86 | 0.72 | 0.84 | -0.35 | -0.06 |
| 5SWD | 65 | 0.0 | 0.0 | 0.0 | 6.7 | 7.5 | 3 | 1 | 1 | 4 | 3 | 0.69 | 0.27 | 0.38 | 0.61 | 0.60 |
| 6C27 | 47 | 9.7 | 12.3 | 11.7 | 9.7 | 11.4 | 5 | 10 | 4 | 4 | 3 | 0.60 | 0.69 | 0.46 | 0.69 | 0.74 |
| 6E8S | 38 | 0.0 | 22.7 | 21.2 | 21.6 | 21.1 | 30 | 29 | 25 | 32 | 33 | 0.26 | -0.07 | 0.34 | -0.09 | -0.05 |
| 6JQ6 | 81 | 33.3 | 42.1 | 32.3 | 32.9 | 32.3 | 7 | 30 | 9 | 9 | 8 | 0.21 | -0.03 | 0.03 | -0.03 | -0.05 |
| 6TFE | 52 | 0.0 | 0.0 | 12.5 | 0.0 | 15.6 | 35 | 30 | 33 | 29 | 29 | 0.38 | -0.04 | 0.37 | -0.01 | -0.25 |
| 6VMY | 148 | 0.0 | 0.0 | 48.0 | 27.0 | 29.8 | 13 | 12 | 14 | 2 | 11 | 0.47 | 0.42 | 0.46 | 0.68 | 0.65 |
| 7D82 | 50 | 17.7 | 14.6 | 9.0 | 14.6 | 17.7 | 10 | 3 | 23 | 25 | 21 | -0.03 | -0.09 | -0.10 | -0.33 | -0.34 |
| 7K16 | 51 | 0.0 | 0.0 | 0.0 | 5.2 | 5.2 | 2 | 6 | 3 | 1 | 1 | 0.73 | 0.25 | 0.65 | 0.66 | 0.66 |
| 7KJU | 75 | 0.0 | 0.0 | 20.5 | 19.8 | 27.0 | 7 | 28 | 13 | 4 | 4 | 0.66 | 0.74 | 0.42 | 0.71 | 0.74 |
| Average |  | **3.3** | 6.2 | 10.2 | 15.9 | 17.0 | **8.1** | 10.8 | 10.8 | 14.8 | 14.4 | **0.64** | 0.47 | 0.54 | 0.19 | 0.14 |

Therefore, for the subsets in Test set III from realistic prediction models, our rsRNASP is always apparently superior to the existing statistical potentials in identifying native structures and in ranking decoy ones.

### Overall performance of rsRNASP on all test sets

#### In identifying native structures

Overall, as shown in Table 1, for all the test sets (test sets I and II with small RNAs or near-native decoys mainly from perturbation methods, and realistic test set III with large RNAs of large RMSD range from various structure prediction models), rsRNASP has apparently the best performance in identifying native structures, since rsRNASP can identifying 145 native structures and 157 native structures within top five ones of lowest energies out of 167 RNA decoy sets. However, such two evaluation numbers are 119 and 148 for RNA3DCNN, 137 and 144 for DFIRE-RNA, 122 and 129 for 3dRNAscore, and 100 and 117 for RASP, respectively. Furthermore, as shown in Figs. 2A and B, rsRNASP can identify 87% native structures and 94% native structures with top five ones with lowest energies for all test sets and even for test set III from various structure prediction models, shown in Figs. 2A and B, such two percentages can reach 76% and 88%. This indicates the apparently better performance of rsRNASP in identifying native structures, especially for the realistic test set III from various structure prediction models.

**Figure 2.**
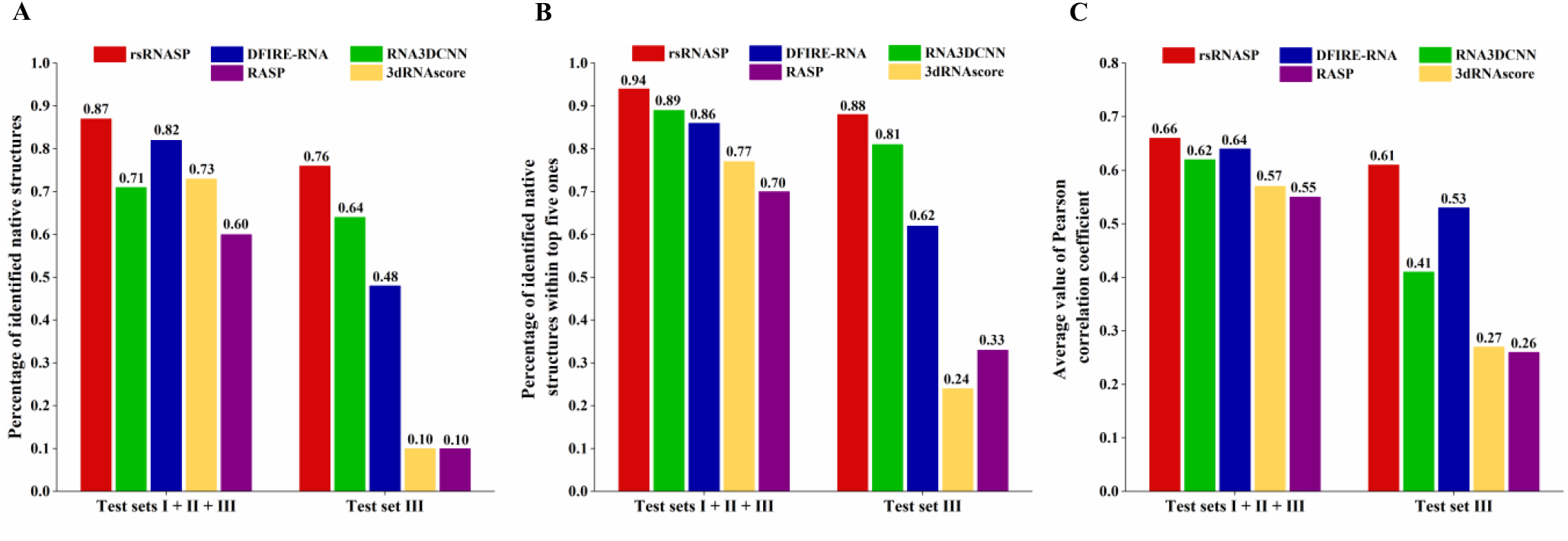
(A) The percentages of the number of identified native structures, (B) the percentages of the numbers of native structures identified within top five ones of lowest energies, and (C) the average values of PCCs on DIs by different statistical potentials for the whole test sets (test sets I+II+III) and test set III, respectively. Here, the PCC values for the whole test sets and for test set III were averaged over the mean values of individual test set and subsets since decoys in a test set or subset were generated with a same method and generally have similar structure features. It is noted that the performance on test set III was explicitly shown because test set III is considered as a realistic test set in which decoy structures were generated from various 3D structure prediction models.

#### In ranking decoy structures

As shown in Fig. 2C, rsRNASP has an overall better performance than other statistical potentials in ranking decoy structures for all test sets since the average PCC values for all test sets are 0.66 for rsRNASP, 0.62 for RNA3DCNN, 0.64 for DFIRE-RNA, 0.57 for 3dRNAscore and 0.55 for RASP, respectively. Such superiority of rsRNASP becomes very apparent for test set III from various 3D structure prediction models, and the PCC value from rsRNASP can still reach ∼0.61 which is visibly larger than those from RNA3DCNN (∼0.41), DFIRE-RNA (∼0.53), 3dRNAscore (∼0.27) and RASP (∼0.26). Therefore, overall, rsRNASP performs better than other statistical potentials in ranking decoy structures, especially for realistic test set III from various structure prediction models.

### Contributions of short- and long-ranged interactions

Since our rsRNASP has apparently better performance for test set III, we examined the individual contributions of short- and long-ranged interactions against RNA-Puzzles_standardized and PM subsets in test set III, as shown in Fig. 3 and Table S9 and S10 in the Supporting Material. The short-ranged contribution performs better than long-ranged one in identifying native structures for both Puzzles_standardized and PM subsets. This indicates that the difference between native structures and their corresponding predicted decoy structures can be more effectively captured by the short-ranged interactions. Furthermore, in ranking the nearest-native structure (on DIs) and ranking decoys (reflected by PCC values on DIs), the long-ranged potential is overall superior to the short-ranged one for Puzzles_standardized subset, while for PM subset, the short-ranged potential has better overall performance in these two metrics. The different performances of the short-ranged and long-ranged interactions against the two subsets from prediction models might come from the different methods for generating the subsets. The decoy structures in PM subset were predicted by the four prediction models with given secondary structures from their 3D ones and consequently their short-ranged interactions can be better captured by the short-ranged potential extracted from native structure ensemble. However, Puzzles_standardized subset was produced by various prediction models possibly without given secondary structures in some cases and may contain mis-folded secondary structures, and thus the long-ranged potential can relatively make more contribution than the short-ranged one in identifying and ranking near-native structures. Certainly, the combination of short-ranged and long-ranged potentials overall performs better than the individual short-ranged or long-ranged potential, reflected by the average values of three metrics (DI of structure with the lowest energy, rank of the nearest-native structure and PCC). Therefore, the overall excellent performance of rsRNASP for different metrics should be attributed to the classification of interaction ranges.

**Figure 3.**
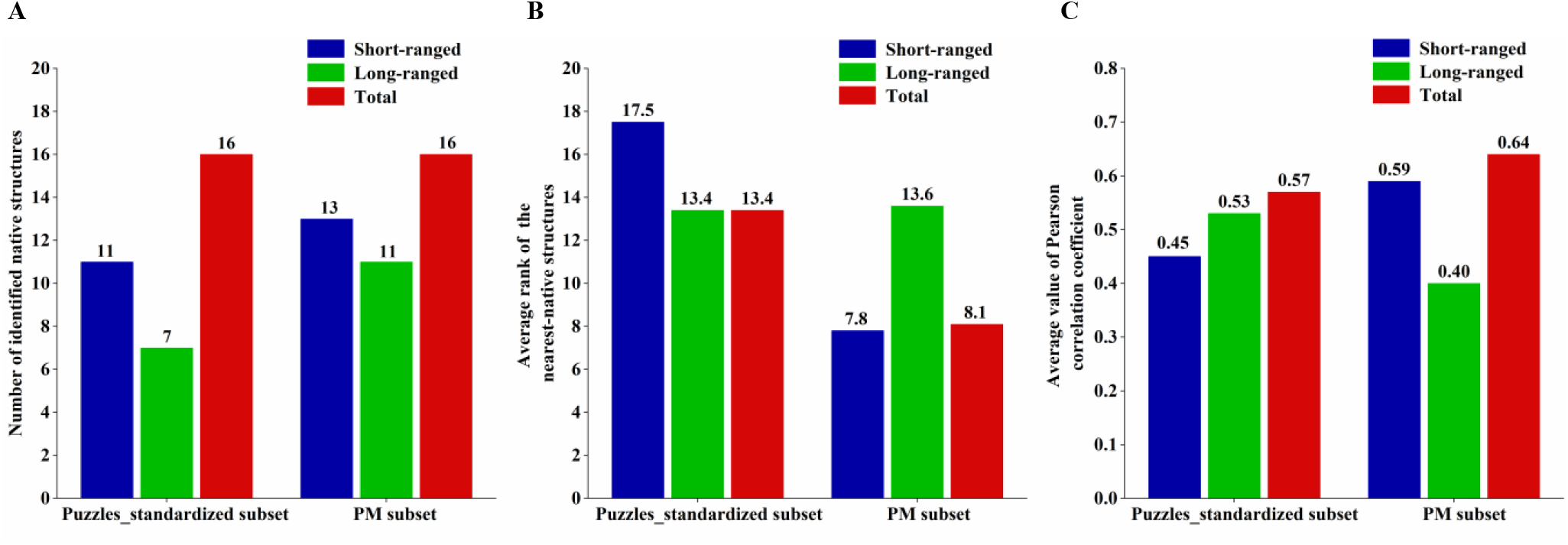
(A) The number of identified native structures, (B) the average rank of the nearest-native structures, and (C) the average value of PCCs on DIs by short-ranged and long-ranged interactions, and by the whole potentials of rsPNASP for Puzzles_standardized and PM subsets in test set III, respectively.

### Ability of capturing physical interactions in RNA

In the following, we analyzed the interactions captured by the short- and long-ranged potentials in rsRNASP, which is responsible for the excellent performance of rsRNASP for the realistic test set III from structure prediction models. As a paradigm, Figs. 4A and B show the short-ranged and long-ranged potentials between N9 atom of adenine (A) and N1 atom of uracil (U), respectively. In the short-ranged potential, the deepest potential well at the distance of ∼7.1 Å is mainly corresponding to the reverse Hoogsteen base-pairing interaction (89), and a weaker potential well at the distance of ∼ 8.9 Å captures the canonical Watson-Crick base-pairing; see Figs. 4C and D. Here, compared with canonical Watson-Crick base-pairing, the reverse Hoogsteen base-pairing mode is more preferred due to the low-frequency Watson-Crick base-pairing in the limited short-ranged residue separation. In addition, another potential well at the distance of ∼5.7 Å represents the intra-loop base interactions separated by two or three residues in the loop region; see Fig. 4E. In the long-ranged potential, as shown in Fig. 4B, there are four local potential wells within the distance of 10 Å. The two deepest ones at the distances of ∼8.9 Å and ∼7.1 Å capture the Watson-Crick and the reverse Hoogsteen base-pairing interactions between A and U, respectively; see in Figs. 4C and D. Besides, the interaction between the nearest diagonal AN9 and UN1 in helix region also appears at ∼7.1 Å; see Fig. 4F. Moreover, the other two potential wells at the distance of ∼4.1 Å and ∼5.0 Å correspond to the different kinds of tertiary base-stacking interactions, including base-stacking between two residues in two adjacent branches of RNA 3D structures (e.g., tetraloop receptor), coaxial-stacking at junction region and base-stacking in triple-helical region; see Figs. 4G-I and Fig. S12 in the Supporting Material.

**Figure 4.**
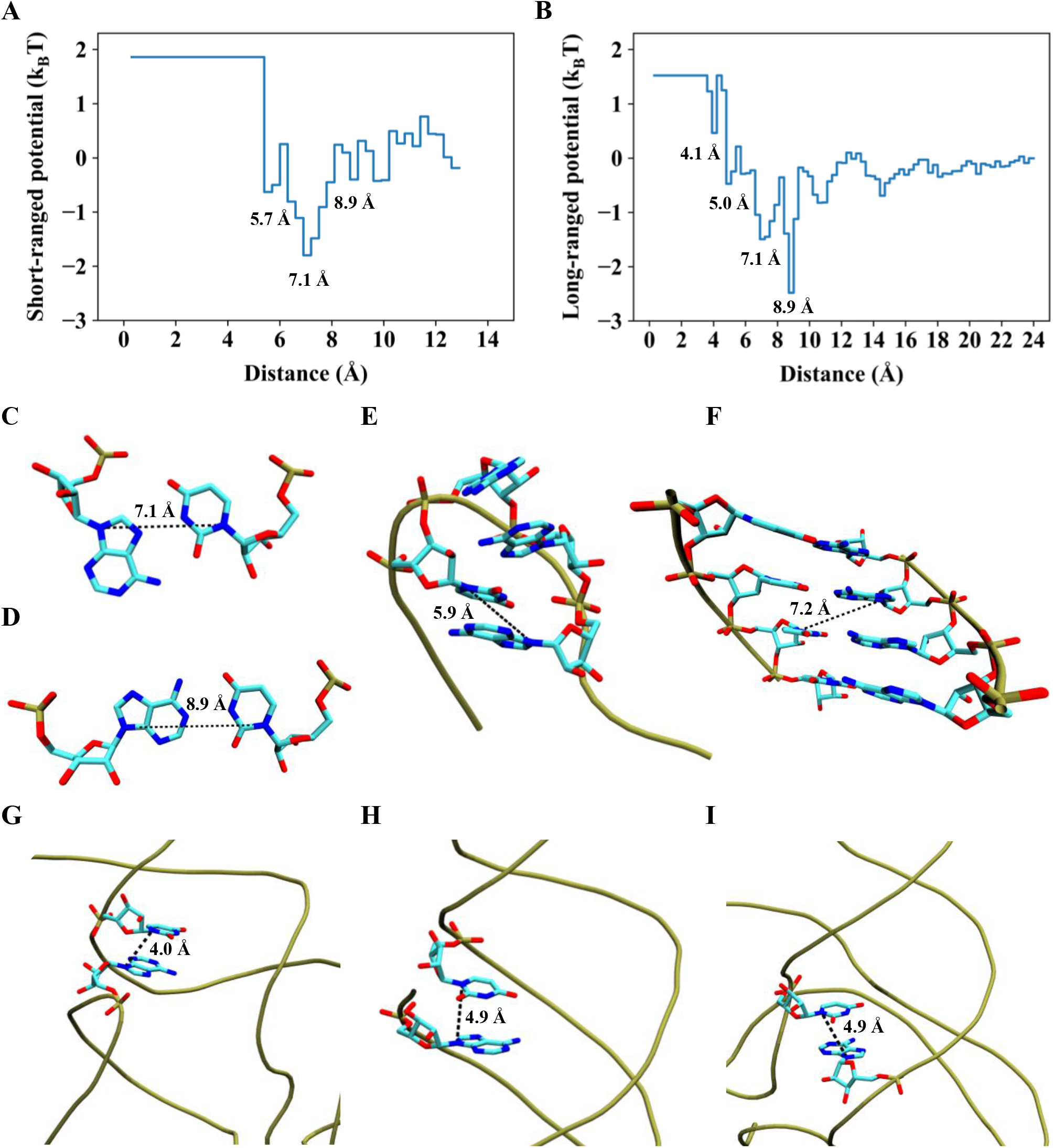
(A) Short-ranged and (B) long-ranged potentials between AN9 and UN1 in rsRNASP. (C) and (D) Representative distances between AN9 and UN1 for reverse Hoogsteen (C) and Watson-Crick (D) base-pairing interactions. (E) Representative distance between AN9 and UN1 for intra-loop base interaction in the short-ranged potential (PDB ID: 4GXY). (F)-(I) Representative distances between AN9 and UN1 for the nearest diagonal base interaction of A-form helix (F), base-stacking interaction of two residues in adjacent branches of RNA 3D structure (G) (PDB ID: 6DME), base-stacking interaction of triple-helical region (H) (PDB ID: 3P22) and coaxial-stacking interaction at junction region (PDB ID: 4QK9) in the long-ranged potential, respectively. Please see Fig. S12 for the complete 3D structures for panels G-I.

As shown above, compared with a potential without distinguishing residue separation (61, 63), the short- and long-ranged potentials can capture more interaction information and especially, the long-ranged potential can capture some kinds of tertiary interactions, which should be responsible for the good performance of rsRNASP for the realistic test set III from structure prediction models.

## Discussion

The most important difference between rsRNASP and other statistical potentials for RNAs is the classification of the short- and long-ranged interactions in rsRNASP. Such classification for interaction ranges may extract more details in the potentials such as reverse Hoogsteen base-pairing and some tertiary base-stacking interactions described above. Furthermore, for extracting short- and long-ranged potentials, we employed the average and the random-walk reference states, respectively. Such hybridization of a database-based reference state and a physical model-based one may bring a complementary effect for a potential (65). Moreover, the very strong local (connectivity-related) interactions within two residue separations were excluded in building rsRNASP, and consequently the nonlocal interactions for stabilizing global/tertiary structures becomes more important in rsRNASP, which would be helpful for evaluating global structures of large RNAs. The above-described treatments involved in rsRNASP should contribute to the apparently higher overall performance of rsRNASP than the existing statistical potentials for the realistic test set III.

Here, we would also like to discuss the performance of the existing top statistical potentials or scoring functions. The excellent performance on test sets I and II and very weak performance on test set III for 3dRNAscore may come from the involvement of local torsional angle potential, which emphasizes the local structure information and consequently under-reflects the long-ranged structure information (63). The relatively good performance on test set II and test set III and weak performance on test set I for RNA3DCNN may be attributed to its training decoy sets containing numerous decoys from MD simulations and Rosetta structure prediction model which may bring a bias for evaluating RNA decoy sets generated by MD method and from structure predictions (67). The overall modestly good performance of DFIRE-RNA for different test sets may be attributed to the balance between short-ranged and long-ranged structure information through using a large bin width (64). The overall worst performance of RASP may come from the lower resolution of atom types and limited structure data in the training set (62).

## CONCLUSIONS

In this work, we have developed a novel knowledge-based potential (rsRNASP) for RNA 3D structure evaluation. rsRNASP was built through distinguishing short- and long-ranged interactions and through using different reference states and training on a non-redundant training native set and a training decoy set. We also built an additional realistic decoy test subset (PM subset) for benchmark test. The extensive tests against the available test sets indicate that, overall, rsRNASP has consistently good performance for test sets I, II and III, especially for the realistic test set III including two subsets from realistic structure prediction models. Furthermore, rsRNASP performs visibly better than existing statistical potentials for the realistic test set III. It is encouraging that our “traditional” rsRNASP is superior to the recently developed RNA3DCNN from 3D convolution neural network.

Of course, rsRNASP (and other “traditional” statistical potentials) can be furtherly improved for more accurate evaluation on RNA 3D structures. First, the reference states can be circumvented through some treatments such as iterative technique since the employed reference states may still deviate largely from the ideal one (52, 55). Second, other geometric parameters such as torsional angle and orientation beyond atom-atom distance can be involved to enhance the description of the relative relationship between atoms or atom groups (29, 55, 63). Third, the classification of interaction ranges can become more subtle and more interaction ranges would bring more weight parameters and consequently more accurate evaluation for structures (55, 71). Fourth, a multi-body (three- or four-body) potential can be supplemented to a pair-wise one which would improve the description on atom-atom distributions (65, 66). Finally, a statistical potential is still limited by the limited RNA native structures in PDB, and it can be improved with the increase of the number of RNA structures deposited in PDB database (55). Additionally, for a statistical potential from training through neural network, more extensive training decoy set built from extensive methods would effectively diminish the training bias of a statistical potential and certainly improve its ability in evaluating RNA 3D structures.

## AVALIABLITY

rsRNASP and the relevant databases are available at website https://github.com/Tan-group/rsRNASP.

## SUPPORTING MATERIAL

Supporting Material is available for this article.

## AUTHOR CONTRIBUTIONS

ZJT, WZ and YLT designed the research. YLT, XW and YZS performed the research. ZJT, YLT and WZ analyzed the data. YLT, YZS, and ZJT wrote the manuscript.

## ACKNOWLEDGMENTS

We are grateful to Profs Shi-Jie Chen (University of Missouri) and Jian Zhang (Nanjing University) for valuable discussions.

This work was supported by grants from the National Science Foundation of China (11774272, 12075171). Parts of the numerical calculations in this work were performed on the super computing system in the Super Computing Center of Wuhan University.

