## Supporting Material for "rsRNASP: A residue-separation-based statistical potential for RNA 3D structure evaluation"

### 1. A detailed description of deriving the statistical potential of rsRNASP

Our present rsRNASP is composed of short- and long-ranged energy functions distinguished by residue separation $k$, which is defined by $k = |m - n|$, where $m$ and $n$ correspond to the observed residue sequence positions of a pair of atoms along an RNA chain.

For extracting information of short-ranged potential, the averaging reference state was used (1). Besides, to avoid the problem of sparse data in short-ranged potential, we employed the method developed by Sippl (2). Hence, the short-ranged potential can be expressed by:

$$\Delta E_{\text{short}}(i,j,r) = k_B T\ln[1 + M_{ij}\sigma] - k_B T\ln\left[1 + M_{ij}\sigma\frac{P_{2<k\le k_0}^{\text{obs}}(i,j,r)}{P_{2<k\le k_0}^{\text{obs}}(r)}\right], \tag{1}$$

where $k_0$ is the residue separation threshold to distinguish between short- and long-ranged interactions. $P_{2<k\le k_0}^{\text{obs}}(i,j,r)$ is the observed probability of the distance between atom pair of atom types $i$ and $j$ within residue separation interval $(2, k_0]$ in distance interval $(r, r + dr]$, and $P_{2<k\le k_0}^{\text{obs}}(r)$ is the summation of $P_{2<k\le k_0}^{\text{obs}}(i,j,r)$ over all kinds of atom pair types. $M_{ij}$ is the total number atom pair of atom types $i$ and $j$ observed within residue separation intervals $(2, k_0]$. $\sigma$ was set to 0.02, as proposed by Sippl (2).

For long-ranged potential, the random-walk-chain reference state was used (3). Thus, according to previous description (4), the probability of reference state $P_{k>k_0}^{\text{ref}}(i,j,r)$ can be given by (4):

$$P_{k>k_0}^{\text{ref}}(i,j,r) = \sum_p P_{k>k_0}^{\text{obs,p}}(i,j,r_c)\left(\frac{r}{r_c}\right)^2\frac{\sum_{n=k_0+1}^{N}\exp(-3r^2/2nl^2)/n^{3/2}}{\sum_{n=k_0+1}^{N}\exp(-3r_c^2/2nl^2)/n^{3/2}}, \tag{2}$$

where $P_{k>k_0}^{\text{obs,p}}(i,j,r_c)$ is the observed probability of the distance between atom pair of atom types $i$ and $j$ within residue separation interval $(k_0, N]$ at distance cutoff $r_c$ for one single native RNA. $N$ is the number of residues in the example structure $p$. $l$ is the Kuhn length, and $l^2$ was set to 310 Å$^2$, as used in our previous work (4). Thus, the long-ranged potential can be given by:

$$\Delta E_{\text{long}}(i,j,r) = -k_B T\ln\frac{P_{k>k_0}^{\text{obs}}(i,j,r)}{\sum_p P_{k>k_0}^{\text{obs,p}}(i,j,r_c)\left(\frac{r}{r_c}\right)^2\frac{\sum_{n=k_0+1}^{N}\exp(-3r^2/2nl^2)/n^{3/2}}{\sum_{n=k_0+1}^{N}\exp(-3r_c^2/2nl^2)/n^{3/2}}}. \tag{3}$$

### 2. Determination of residue separation threshold $k_0$

The residue separation threshold $k_0$ was introduced to distinguish short- and long-ranged interactions. A long-ranged potential extracted from training set is aimed to describe non-local interactions in RNAs, including base pairing and crucial tertiary interactions. A short-ranged potential in RNAs is designated to represent the features of RNA local loop regions as accurate as possible. Hence, $k_0$ describes the residue separation boundary between intra-loop interactions and those beyond in RNA structures. With x3DNA-DSSR (5), the secondary structures of 191 native RNAs in training set were parsed into dot-bracket forms, and we can obtain the frequency distribution of numbers of consecutive dots through statistical analyses, namely number of residues occupied by local loop regions. As shown in Fig. S1, we found that a sharp decrease appears after the residue separation $k$ of 4. Thus, the residue separation threshold $k_0$ was taken as 4 to distinguish short- and long-ranged interactions in the rsRNASP.

**3. Determination of distance cutoffs for short- and long-ranged potentials**

According to the frequency distribution of spatially long-ranged atom pairs ($k > k_0$=4) for 191 native RNAs in training set, we found that the curve become saturated and afterwards declined after ~24 Å; see Fig. S2. Thus, the distance cutoff $r_c$ for long-ranged interactions was set to 24 Å.

Similarly, the frequency distribution of short-ranged atom pairs ($k <= k_0$=4) for native RNAs in training set showed a peak at ~14 Å; see Fig. S3. Thus, the distance cutoff $r_c$ for short-ranged interactions was set to 13 Å, for excluding possible helix structure information in extracting the information of RNA local loop regions.

**4. Optimization of weight $\omega$ between short- and long-ranged potentials**

Different from strong short-ranged interactions limited by much less residue separation and shorter distance cutoff, for an atom, the number of its atom pairs of long-ranged interactions per atom (or per nucleotide) could strongly depend on RNA length $N$ (in nt). Thus, at first, a function related to RNA length $N$ (in nt) is required to normalize the energy contribution from long-ranged interactions. The relation between the number of long-ranged atom pairs within 24 Å scaled by $N$ and RNA length for 191 native RNAs in training native set was shown in Fig. S4. According to Fig. S4, for convenience and simplicity, we employed a fitted function in our rsRNASP:

$$f(N) = \frac{-2685}{(N+16)^{0.5}} + 542, \quad (4)$$

where $N$ is the nucleotide length of an RNA structure. Therefore, the weight $w$ in the rsRNASP can be furtherly given by:

$$w = w_0/f(N). \quad (5)$$

Here, $w_0$ is a constant weight required to be determined. In our rsRNASP, $w_0$ was determined by the training decoy set which was built from the RNA 3D structure prediction models (including FARFAR2 (6), RNAComposer (7), SimRNA (8) and 3dRNA v2.0 (9)) and includes 35 single-stranded RNAs with about 40 decoy structures for each RNA. Specifically, two metrics, including the number of native structures identified within top five of the lowest energies and Pearson correlation coefficient (PCC) values on DIs (10), can be obtained under different weight by leave-one-out way (11). As shown in Fig. S5, the choice of the value of $w_0$=16 can maximize these two metrics, i.e., $w_0$=16 is the most optimized value for the the training decoy set of 35 single-stranded RNAs with about 40 decoy structures for each RNA from the four structure prediction models (5-8).

Table S1. PDB IDs of 191 RNAs in our training native set[a].

| | | | | | | | | | | | | |
|---|---|---|---|---|---|---|---|---|---|---|---|---|
| 1CSL | 1D4R | 1DUH | 1DUQ | 1ET4 | 1EVV | 1F1T | 1F27 | 1FIR | 1FUF | 1I9X | 1KD5 | 1KFO |
| 1KH6 | 1KXK | 1L2X | 1MHK | 1NBS | 1NTA | 1NUV | 1Q96 | 1QBP | 1QCU | 1RNA | 1SDR | 1T0E |
| 1U9S | 1XJR | 1Y26 | 1Y27 | 1YFG | 1YLS | 1Z7F | 1ZCI | 205D | 255D | 280D | 283D | 2A64 |
| 2ET8 | 2G92 | 2GDI | 2H1M | 2IL9 | 2JLT | 2O3Y | 2OE8 | 2OEU | 2OIU | 2P7E | 2Q1R | 2QUS |
| 2QWY | 2R1S | 2YGH | 2ZY6 | 353D | 359D | 361D | 364D | 387D | 397D | 3BNN | 3CGS | 3CJZ |
| 3CZW | **3D2V** | 3DIL | 3E5C | 3GM7 | 3GS5 | 3IBK | 3IGI | 3K1V | 3LOA | 3MEI | 3MJA | 3ND3 |
| 3NPQ | 3OXE | 3P22 | 3P59 | 3PDR | **3Q3Z** | **3R4F** | **3RG5** | 3SJ2 | **3SKL** | **3SUX** | 406D | 413D |
| 422D | 433D | 4E48 | 4E5C | **4ENC** | **4FRN** | 4GXY | 4J50 | **4JF2** | **4JRC** | 4JRD | 4JRT | **4K27** |
| **4KQY** | 4KYY | **4LVW** | 4MCF | 4NFQ | 4NLF | 4O41 | 4OQU | 4P3T | 4P95 | 4P97 | 4PCJ | 4PHY |
| **4PLX** | **4PQV** | 4QJD | 4QK9 | 4QLM | 4R4V | 4RBQ | 4RBY | 4RGE | 4RUM | 4RZD | 4TS2 | **4WFL** |
| 4XWF | **4Y1M** | **4YAZ** | **4ZNP** | 5AY2 | 5BJO | **5BTM** | 5BTP | 5C5W | 5CNR | 5EW4 | 5G4T | 5KPY |
| 5KTJ | 5KVJ | 5L4O | **5LYS** | **5M0H** | **5ML7** | 5MWI | 5NDI | 5NWQ | 5NXT | **5OB3** | 5T83 | **5U3G** |
| 5UNE | 5UZ6 | 5V1K | 5V3F | 5VJ9 | 5XWG | 5Z1I | 6C63 | 6CB3 | 6CK5 | 6CU1 | 6D3P | **6DLR** |
| **6DME** | 6DN2 | **6DVK** | 6E1S | 6E7L | **6E8U** | 6FZ0 | **6H0R** | 6HC5 | 6HU6 | 6IA2 | 6JQ5 | **6JXM** |
| **6MJ0** | **6N2V** | **6N5P** | 6OL3 | **6P2H** | 6PMO | 6QN3 | 6R47 | 6UFG | | | | |

[a] PDB IDs of the 35 RNAs in the training decoy set are marked in bold.

Table S2. The average ES and PCC values calculated by different statistical potentials for test set I.

| Test set I | Enrichment score | | | | | Pearson correlation coefficient | | | | |
|---|---|---|---|---|---|---|---|---|---|---|
| | rsRNASP | RNA3DCNN | DFIRE-RNA | 3dRNAscore | RASP | rsRNASP | RNA3DCNN | DFIRE-RNA | 3dRNAscore | RASP |
| Average value (DIs) | 8.6 | 8.9 | 8.7 | **9.0** | **9.0** | 0.80 | 0.84 | 0.81 | 0.82 | **0.86** |
| Average value (RMSDs) | 8.4 | 8.8 | 8.5 | **8.9** | 8.8 | 0.83 | 0.88 | 0.83 | 0.87 | **0.89** |

Table S3. The ES and PCC values on DIs calculated by different statistical potentials for test set II.

| | | | Enrichment score (DI) | | | | | Pearson correlation coefficient (DI) | | | | |
|---|---|---|---|---|---|---|---|---|---|---|---|---|
| Decoy sets | RNA | length | rsRNASP | RNA3DCNN | DFIRE-RNA | 3dRNAscore | RASP | rsRNASP | RNA3DCNN | DFIRE-RNA | 3dRNAscore | RASP |
| Test set II_MD | 1duq | 26 | 7.8 | 7.3 | 6.9 | 8.3 | 7.7 | 0.83 | 0.69 | 0.85 | 0.74 | 0.74 |
| | 1f27 | 30 | 7.1 | 4.2 | 6.9 | 8.1 | 6.5 | 0.87 | 0.71 | 0.89 | 0.71 | 0.71 |
| | 1msy | 27 | 6.3 | 7.3 | 6.5 | 7.8 | 5.7 | 0.85 | 0.89 | 0.84 | 0.90 | 0.88 |
| | 1nuj | 24 | 6.5 | 8.6 | 7.3 | 7.6 | 5.2 | 0.77 | 0.67 | 0.78 | 0.73 | 0.69 |
| | 434d | 14 | 7.6 | 8.5 | 7.8 | 7.6 | 7.0 | 0.72 | 0.68 | 0.74 | 0.68 | 0.67 |
| Average | | | 7.1 | 7.2 | 7.1 | **7.9** | 6.4 | 0.81 | 0.73 | **0.82** | 0.75 | 0.74 |
| Test set II_NM | 1duq | 26 | 6.3 | 6.1 | 7.1 | 7.1 | 5.9 | 0.90 | 0.86 | 0.91 | 0.91 | 0.85 |
| | 1esy | 19 | 4.1 | 7.5 | 5.9 | 5.5 | 5.1 | 0.92 | 0.87 | 0.94 | 0.92 | 0.87 |
| | 1f27 | 30 | 4.7 | 6.3 | 5.3 | 6.3 | 3.7 | 0.92 | 0.91 | 0.94 | 0.90 | 0.87 |
| | 1i9v | 76 | 4.5 | 4.3 | 3.1 | 6.3 | 5.7 | 0.90 | 0.90 | 0.88 | 0.92 | 0.91 |
| | 1kka | 17 | 4.7 | 3.7 | 6.1 | 6.1 | 4.1 | 0.84 | 0.85 | 0.84 | 0.84 | 0.76 |
| | 1msy | 27 | 5.5 | 7.1 | 5.3 | 6.5 | 2.9 | 0.91 | 0.88 | 0.93 | 0.94 | 0.92 |
| | 1nuj | 24 | 6.3 | 7.5 | 6.5 | 7.1 | 6.1 | 0.92 | 0.85 | 0.94 | 0.91 | 0.89 |
| | 1qwa | 21 | 2.9 | 5.1 | 2.6 | 3.9 | 2.6 | 0.77 | 0.79 | 0.79 | 0.80 | 0.68 |
| | 1x9k | 62 | 3.7 | 5.2 | 1.9 | 6.2 | 5.4 | 0.89 | 0.91 | 0.90 | 0.90 | 0.86 |
| | 1xjr | 46 | 8.3 | 6.7 | 7.7 | 7.9 | 8.3 | 0.94 | 0.91 | 0.94 | 0.94 | 0.95 |
| | 1ykq | 49 | 6.6 | 5.2 | 3.1 | 5.2 | 3.7 | 0.90 | 0.88 | 0.91 | 0.90 | 0.88 |
| | 1zih | 12 | 6.7 | 5.9 | 7.3 | 7.3 | 5.9 | 0.92 | 0.92 | 0.94 | 0.93 | 0.89 |
| | 28sp | 28 | 5.5 | 6.9 | 6.1 | 5.9 | 6.7 | 0.85 | 0.89 | 0.92 | 0.93 | 0.92 |
| | 2f88 | 34 | 6.0 | 6.2 | 4.9 | 6.2 | 4.9 | 0.92 | 0.92 | 0.93 | 0.92 | 0.91 |
| | 434d | 14 | 7.9 | 7.7 | 7.9 | 8.4 | 7.9 | 0.92 | 0.90 | 0.92 | 0.93 | 0.90 |
| Average | | | 5.6 | 6.1 | 5.4 | **6.4** | 5.3 | 0.89 | 0.88 | **0.91** | 0.90 | 0.87 |
| Test set II_FARNA | 157d | 24 | 4.4 | 3.0 | 3.2 | 3.0 | 3.8 | 0.58 | 0.58 | 0.50 | 0.62 | 0.68 |
| | 1a4d | 41 | 1.6 | 3.4 | 1.4 | 2.2 | 3.0 | 0.09 | 0.24 | 0.10 | 0.19 | 0.38 |
| | 1csl | 28 | 2.0 | 3.0 | 1.6 | 2.6 | 2.0 | 0.35 | 0.59 | 0.16 | 0.45 | 0.44 |
| | 1dqf | 19 | 3.6 | 5.6 | 2.2 | 3.8 | 3.0 | 0.53 | 0.68 | 0.41 | 0.64 | 0.48 |
| | 1esy | 19 | 1.8 | 4.0 | 2.4 | 3.8 | 4.8 | 0.27 | 0.59 | 0.39 | 0.63 | 0.64 |
| | 1i9x | 26 | 5.8 | 3.2 | 3.0 | 3.6 | 2.6 | 0.68 | 0.41 | 0.43 | 0.51 | 0.36 |
| | 1j6s | 24 | 3.0 | 1.2 | 2.6 | 1.2 | 2.2 | 0.45 | 0.19 | 0.54 | 0.30 | 0.52 |
| | 1kd5 | 22 | 0.8 | 4.4 | 0.6 | 2.6 | 0.8 | 0.12 | 0.53 | 0.12 | 0.47 | 0.30 |
| | 1kka | 17 | 0.0 | 3.8 | 0.2 | 2.0 | 1.0 | -0.18 | 0.66 | -0.12 | 0.28 | 0.08 |
| | 1l2x | 27 | 2.0 | 0.2 | 3.8 | 0.8 | 1.2 | 0.35 | -0.49 | 0.62 | -0.14 | -0.13 |
| | 1mhk | 32 | 0.8 | 2.0 | 1.0 | 1.2 | 0.4 | 0.14 | 0.26 | 0.15 | 0.30 | 0.18 |
| | 1q9a | 27 | 0.6 | 3.8 | 0.6 | 3.0 | 1.0 | 0.06 | 0.48 | 0.03 | 0.46 | 0.16 |
| | 1qwa | 21 | 0.2 | 3.6 | 0.2 | 1.2 | 0.2 | 0.03 | 0.44 | -0.08 | 0.28 | 0.26 |
| | 1xjr | 46 | 3.4 | 2.4 | 3.2 | 3.0 | 3.2 | 0.40 | 0.56 | 0.37 | 0.50 | 0.56 |
| | 1zih | 12 | 4.8 | 10.0 | 6.8 | 6.8 | 7.2 | 0.60 | 0.77 | 0.62 | 0.62 | 0.47 |
| | 255d | 24 | 0.8 | 3.6 | 0.4 | 2.2 | 0.6 | -0.26 | 0.71 | -0.25 | 0.43 | -0.14 |
| | 283d | 24 | 1.0 | 0.2 | 1.0 | 1.4 | 0.6 | 0.12 | 0.11 | 0.24 | 0.21 | 0.31 |
| | 28sp | 28 | 2.4 | 4.0 | 3.2 | 4.4 | 4.0 | 0.28 | 0.62 | 0.06 | 0.57 | 0.49 |
| | 2a43 | 26 | 1.2 | 3.4 | 2.6 | 3.0 | 2.6 | 0.14 | 0.25 | 0.47 | 0.31 | 0.27 |
| | 2f88 | 34 | 1.8 | 4.0 | 1.4 | 3.4 | 1.6 | 0.22 | 0.40 | 0.13 | 0.47 | 0.21 |
| Average | | | 2.1 | **3.4** | 2.1 | 2.8 | 2.3 | 0.25 | **0.43** | 0.24 | 0.40 | 0.33 |

Table S4. The ES and PCC values on RMSDs calculated by different statistical potentials for test set II.

| Decoy sets | RNA | length | Enrichment score (RMSD) | | | | | Pearson correlation coefficient (RMSD) | | | | |
|---|---|---|---|---|---|---|---|---|---|---|---|---|
| | | | rsRNASP | RNA3DCNN | DFIRE-RNA | 3dRNAscore | RASP | rsRNASP | RNA3DCNN | DFIRE-RNA | 3dRNAscore | RASP |
| Test set II_MD | 1duq | 26 | 7.9 | 7.3 | 6.9 | 8.3 | 7.7 | 0.89 | 0.67 | 0.90 | 0.78 | 0.73 |
| | 1f27 | 30 | 7.0 | 4.2 | 6.9 | 8.1 | 6.6 | 0.88 | 0.66 | 0.89 | 0.68 | 0.66 |
| | 1msy | 27 | 6.4 | 7.3 | 6.7 | 7.9 | 5.7 | 0.83 | 0.87 | 0.81 | 0.89 | 0.87 |
| | 1nuj | 24 | 6.4 | 8.6 | 7.3 | 7.6 | 5.2 | 0.93 | 0.66 | 0.92 | 0.79 | 0.69 |
| | 434d | 14 | 7.6 | 8.5 | 7.8 | 7.7 | 7.0 | 0.89 | 0.77 | 0.88 | 0.81 | 0.78 |
| Average | | | 7.1 | 7.2 | 7.1 | **7.9** | 6.4 | **0.88** | 0.73 | **0.88** | 0.79 | 0.75 |
| Test set II_NM | 1duq | 26 | 6.7 | 6.1 | 7.1 | 7.3 | 5.9 | 0.90 | 0.84 | 0.90 | 0.89 | 0.84 |
| | 1esy | 19 | 3.9 | 7.5 | 5.3 | 4.9 | 4.7 | 0.93 | 0.83 | 0.94 | 0.91 | 0.85 |
| | 1f27 | 30 | 4.9 | 6.1 | 5.3 | 6.3 | 3.9 | 0.91 | 0.89 | 0.93 | 0.87 | 0.83 |
| | 1i9v | 76 | 4.5 | 4.3 | 3.1 | 6.3 | 5.7 | 0.90 | 0.90 | 0.89 | 0.92 | 0.91 |
| | 1kka | 17 | 4.7 | 3.5 | 6.1 | 6.1 | 4.3 | 0.91 | 0.90 | 0.91 | 0.90 | 0.80 |
| | 1msy | 27 | 5.7 | 7.1 | 5.1 | 6.3 | 2.4 | 0.91 | 0.88 | 0.93 | 0.94 | 0.91 |
| | 1nuj | 24 | 6.1 | 7.1 | 6.5 | 7.1 | 6.1 | 0.91 | 0.85 | 0.94 | 0.90 | 0.88 |
| | 1qwa | 21 | 3.1 | 4.9 | 2.2 | 3.9 | 2.2 | 0.92 | 0.90 | 0.91 | 0.93 | 0.78 |
| | 1x9k | 62 | 3.7 | 5.4 | 1.9 | 5.8 | 5.4 | 0.88 | 0.90 | 0.87 | 0.88 | 0.86 |
| | 1xjr | 46 | 8.3 | 6.7 | 7.5 | 7.9 | 8.3 | 0.94 | 0.91 | 0.93 | 0.93 | 0.95 |
| | 1ykq | 49 | 6.4 | 5.4 | 3.3 | 5.2 | 3.7 | 0.89 | 0.86 | 0.90 | 0.86 | 0.86 |
| | 1zih | 12 | 6.5 | 5.5 | 7.3 | 7.3 | 5.9 | 0.92 | 0.91 | 0.94 | 0.93 | 0.87 |
| | 28sp | 28 | 5.7 | 6.9 | 6.3 | 5.9 | 6.7 | 0.86 | 0.86 | 0.92 | 0.92 | 0.91 |
| | 2f88 | 34 | 5.7 | 6.2 | 4.7 | 6.2 | 4.9 | 0.93 | 0.91 | 0.92 | 0.92 | 0.91 |
| | 434d | 14 | 7.7 | 7.9 | 7.9 | 8.1 | 7.7 | 0.94 | 0.90 | 0.93 | 0.93 | 0.90 |
| Average | | | 5.6 | 6.0 | 5.3 | **6.3** | 5.2 | 0.91 | 0.88 | **0.92** | 0.91 | 0.87 |
| Test set II_FARNA | 157d | 24 | 3.6 | 2.8 | 2.2 | 2.6 | 3.2 | 0.59 | 0.42 | 0.46 | 0.50 | 0.62 |
| | 1a4d | 41 | 1.2 | 3.4 | 1.2 | 2.0 | 2.4 | 0.06 | 0.20 | 0.08 | 0.15 | 0.33 |
| | 1csl | 28 | 1.6 | 2.0 | 0.8 | 1.8 | 1.2 | 0.33 | 0.51 | 0.10 | 0.37 | 0.35 |
| | 1dqf | 19 | 4.0 | 5.0 | 1.8 | 3.8 | 2.8 | 0.46 | 0.67 | 0.25 | 0.57 | 0.32 |
| | 1esy | 19 | 1.6 | 4.2 | 2.6 | 3.0 | 5.0 | 0.26 | 0.62 | 0.37 | 0.62 | 0.67 |
| | 1i9x | 26 | 5.2 | 2.4 | 2.4 | 2.4 | 2.0 | 0.67 | 0.29 | 0.32 | 0.38 | 0.21 |
| | 1j6s | 24 | 3.4 | 1.0 | 2.8 | 0.8 | 1.4 | 0.54 | -0.07 | 0.44 | -0.04 | 0.14 |
| | 1kd5 | 22 | 1.0 | 3.6 | 0.6 | 2.8 | 0.8 | 0.01 | 0.44 | -0.03 | 0.33 | 0.11 |
| | 1kka | 17 | 0.0 | 3.6 | 0.0 | 2.2 | 0.6 | -0.49 | 0.64 | -0.36 | 0.11 | -0.05 |
| | 1l2x | 27 | 2.2 | 0.2 | 2.6 | 0.4 | 1.0 | 0.39 | -0.54 | 0.64 | -0.18 | -0.20 |
| | 1mhk | 32 | 0.8 | 1.8 | 0.6 | 1.2 | 1.0 | 0.04 | 0.25 | 0.02 | 0.16 | 0.12 |
| | 1q9a | 27 | 0.6 | 3.2 | 0.4 | 3.0 | 0.8 | -0.04 | 0.58 | -0.05 | 0.49 | 0.15 |
| | 1qwa | 21 | 0.2 | 3.8 | 0.2 | 2.0 | 0.4 | -0.19 | 0.73 | -0.26 | 0.40 | 0.21 |
| | 1xjr | 46 | 2.8 | 2.0 | 2.4 | 2.4 | 2.2 | 0.38 | 0.49 | 0.31 | 0.42 | 0.45 |
| | 1zih | 12 | 4.4 | 9.0 | 5.0 | 4.8 | 4.8 | 0.47 | 0.69 | 0.48 | 0.44 | 0.37 |
| | 255d | 24 | 0.8 | 3.6 | 0.4 | 2.0 | 0.6 | -0.31 | 0.67 | -0.34 | 0.37 | -0.23 |
| | 283d | 24 | 1.0 | 0.4 | 1.0 | 1.4 | 0.8 | 0.08 | 0.08 | 0.20 | 0.17 | 0.27 |
| | 28sp | 28 | 1.6 | 3.0 | 2.0 | 3.0 | 3.0 | 0.24 | 0.63 | 0.05 | 0.56 | 0.47 |
| | 2a43 | 26 | 0.8 | 3.0 | 1.4 | 2.0 | 1.4 | 0.13 | 0.22 | 0.45 | 0.24 | 0.14 |
| | 2f88 | 34 | 2.0 | 4.0 | 1.4 | 2.8 | 2.2 | 0.18 | 0.44 | 0.08 | 0.48 | 0.21 |
| Average | | | 1.9 | **3.1** | 1.6 | 2.3 | 1.9 | 0.19 | **0.40** | 0.16 | 0.33 | 0.23 |

Table S5. The RMSDs of structures with the lowest energy and Pearson correlation coefficients between energies and RMSDs of decoy structures calculated by different statistical potentials for Puzzles_standardized subset in test set III.

| | | RMSD of structure with the lowest energy | | | | | Pearson correlation coefficient (RMSD) | | | | |
|---|---|---|---|---|---|---|---|---|---|---|---|
| RNA | length | rsRNASP | RNA3DCNN | DFIRE-RNA | 3dRNAscore | RASP | rsRNASP | RNA3DCNN | DFIRE-RNA | 3dRNAscore | RASP |
| rp01 | 46 | 0.00 | 0.00 | 0.00 | 5.71 | 5.71 | 0.57 | 0.54 | 0.53 | 0.22 | 0.25 |
| rp02 | 100 | 0.00 | 2.30 | 3.66 | 3.66 | 3.66 | 0.59 | 0.40 | 0.28 | 0.20 | 0.38 |
| rp03 | 84 | 0.00 | 14.25 | 0.00 | 14.25 | 14.25 | 0.53 | 0.43 | 0.45 | 0.26 | 0.29 |
| rp04 | 126 | 4.51 | 4.51 | 4.12 | 4.21 | 4.20 | 0.36 | 0.38 | 0.44 | 0.41 | 0.48 |
| rp05 | 188 | 0.00 | 0.00 | 0.00 | 0.00 | 0.00 | 0.43 | 0.65 | 0.62 | 0.73 | 0.73 |
| rp06 | 168 | 14.11 | 0.00 | 14.11 | 28.96 | 30.90 | 0.73 | 0.32 | 0.62 | 0.20 | 0.05 |
| rp07 | 185 | 24.57 | 24.89 | 24.89 | 26.14 | 24.89 | 0.52 | 0.16 | 0.50 | 0.09 | 0.11 |
| rp08 | 96 | 0.00 | 0.00 | 0.00 | 0.00 | 10.65 | 0.74 | 0.59 | 0.74 | 0.41 | 0.66 |
| rp09 | 71 | 6.39 | 6.04 | 6.38 | 6.39 | 6.04 | 0.85 | 0.61 | 0.91 | 0.75 | 0.80 |
| rp10 | 171 | 0.00 | 10.38 | 10.27 | 10.27 | 9.33 | 0.88 | 0.73 | 0.71 | 0.47 | 0.63 |
| rp11 | 57 | 11.54 | 11.54 | 6.17 | 6.17 | 11.54 | -0.23 | 0.22 | -0.27 | -0.11 | -0.07 |
| rp12 | 125 | 13.04 | 0.00 | 15.87 | 13.14 | 13.14 | 0.79 | 0.34 | 0.72 | 0.67 | 0.66 |
| rp13 | 71 | 0.00 | 0.00 | 0.00 | 14.84 | 8.38 | 0.83 | 0.07 | 0.77 | 0.39 | 0.52 |
| rp14_bound | 61 | 0.00 | 0.00 | 12.88 | 5.98 | 7.56 | 0.31 | 0.57 | 0.23 | 0.48 | 0.45 |
| rp14_free | 61 | 0.00 | 0.00 | 6.90 | 11.41 | 11.41 | 0.54 | 0.02 | 0.49 | 0.11 | 0.24 |
| rp15 | 68 | 0.00 | 0.00 | 15.46 | 19.78 | 16.27 | 0.50 | 0.21 | 0.57 | 0.39 | 0.38 |
| rp17 | 62 | 0.00 | 0.00 | 0.00 | 8.64 | 19.07 | 0.39 | -0.02 | 0.40 | 0.23 | 0.16 |
| rp18 | 71 | 0.00 | 0.00 | 0.00 | 16.35 | 12.49 | 0.58 | -0.15 | 0.54 | -0.01 | -0.01 |
| rp19 | 62 | 0.00 | 16.49 | 0.00 | 16.49 | 16.49 | 0.39 | 0.22 | 0.22 | -0.04 | -0.08 |
| rp20 | 68 | 0.00 | 0.00 | 0.00 | 16.24 | 16.41 | 0.40 | 0.26 | 0.41 | -0.02 | -0.23 |
| rp21 | 41 | 0.00 | 4.82 | 5.51 | 18.03 | 18.03 | 0.40 | 0.13 | 0.47 | 0.00 | 0.05 |
| rp24 | 112 | 0.00 | 0.00 | 0.00 | 19.06 | 0.00 | 0.49 | 0.63 | 0.27 | 0.49 | 0.57 |
| Average | | **3.37** | 4.33 | 5.74 | 12.08 | 11.84 | **0.53** | 0.33 | 0.48 | 0.29 | 0.32 |

Table S6. The DIs of structures with the lowest energy and Pearson correlation coefficients between energies and DIs of decoy structures calculated by different statistical potentials for Puzzles_normalized subset in test set III.

| | | DI of structure with the lowest energy | | | | | Pearson correlation coefficient (DI) | | | | |
|---|---|---|---|---|---|---|---|---|---|---|---|
| RNA | length | rsRNASP | RNA3DCNN | DFIRE-RNA | 3dRNAscore | RASP | rsRNASP | RNA3DCNN | DFIRE-RNA | 3dRNAscore | RASP |
| rp01 | 46 | 0.0 | 0.0 | 0.0 | 6.5 | 6.5 | 0.50 | 0.56 | 0.48 | 0.19 | 0.22 |
| rp02 | 100 | 0.0 | 2.8 | 4.3 | 4.3 | 4.3 | 0.65 | 0.44 | 0.36 | 0.29 | 0.42 |
| rp03 | 84 | 0.0 | 21.6 | 0.0 | 21.6 | 21.6 | 0.60 | 0.47 | 0.55 | 0.44 | 0.45 |
| rp04 | 126 | 5.0 | 5.1 | 4.7 | 4.8 | 4.8 | 0.49 | 0.51 | 0.58 | 0.56 | 0.61 |
| rp05 | 188 | 0.0 | 0.0 | 0.0 | 0.0 | 0.0 | 0.44 | 0.65 | 0.63 | 0.76 | 0.76 |
| rp06 | 168 | 19.2 | 0.0 | 19.2 | 40.5 | 45.0 | 0.78 | 0.32 | 0.69 | 0.22 | 0.07 |
| rp07 | 185 | 33.2 | 31.6 | 31.6 | 34.1 | 31.6 | 0.29 | 0.13 | 0.28 | 0.12 | 0.13 |
| rp08 | 96 | 0.0 | 0.0 | 0.0 | 0.0 | 12.4 | 0.70 | 0.47 | 0.74 | 0.34 | 0.64 |
| rp10 | 171 | 0.0 | 12.7 | 12.5 | 12.5 | 12.5 | 0.94 | 0.81 | 0.78 | 0.53 | 0.70 |
| rp12 | 125 | 17.8 | 0.0 | 23.1 | 18.1 | 18.1 | 0.63 | 0.38 | 0.47 | 0.45 | 0.42 |
| rp13 | 71 | 0.0 | 0.0 | 0.0 | 26.5 | 11.1 | 0.83 | 0.31 | 0.90 | 0.84 | 0.87 |
| rp14_bound | 61 | 0.0 | 0.0 | 16.0 | 7.8 | 9.5 | 0.43 | 0.62 | 0.28 | 0.55 | 0.52 |
| rp14_free | 61 | 7.8 | 0.0 | 7.8 | 13.9 | 13.9 | 0.66 | 0.17 | 0.48 | 0.23 | 0.35 |
| rp15 | 68 | 0.0 | 0.0 | 21.7 | 30.4 | 25.0 | 0.54 | 0.12 | 0.54 | 0.38 | 0.39 |
| rp17 | 62 | 0.0 | 0.0 | 0.0 | 11.6 | 25.0 | 0.72 | 0.31 | 0.71 | 0.67 | 0.59 |
| rp19 | 71 | 0.0 | 0.0 | 0.0 | 24.6 | 18.5 | 0.66 | -0.13 | 0.58 | 0.03 | 0.08 |
| rp18 | 62 | 0.0 | 23.0 | 0.0 | 23.0 | 23.0 | 0.23 | 0.13 | 0.00 | -0.22 | -0.23 |
| rp21 | 41 | 0.0 | 0.0 | 0.0 | 34.2 | 34.2 | 0.13 | -0.16 | 0.05 | -0.28 | -0.38 |
| Average | | **4.6** | 5.4 | 7.8 | 17.5 | 17.6 | **0.57** | 0.34 | 0.51 | 0.34 | 0.37 |

Table S7. The RMSDs of structures with the lowest energy and Pearson correlation coefficients between energies and RMSDs of decoy structures calculated by different statistical potentials for Puzzles_normalized subset in test set III.

| | | RMSD of structure with the lowest energy | | | | | Pearson correlation coefficient (RMSD) | | | | |
|---|---|---|---|---|---|---|---|---|---|---|---|
| RNA | length | rsRNASP | RNA3DCNN | DFIRE-RNA | 3dRNAscore | RASP | rsRNASP | RNA3DCNN | DFIRE-RNA | 3dRNAscore | RASP |
| rp01 | 46 | 0.0 | 0.0 | 0.0 | 5.7 | 5.7 | 0.57 | 0.54 | 0.53 | 0.22 | 0.25 |
| rp02 | 100 | 0.0 | 2.3 | 3.7 | 3.7 | 3.7 | 0.59 | 0.40 | 0.28 | 0.20 | 0.38 |
| rp03 | 84 | 0.0 | 14.2 | 0.0 | 14.2 | 14.2 | 0.53 | 0.43 | 0.45 | 0.26 | 0.29 |
| rp04 | 126 | 4.5 | 4.5 | 4.1 | 4.2 | 4.2 | 0.36 | 0.38 | 0.44 | 0.41 | 0.48 |
| rp05 | 188 | 0.0 | 0.0 | 0.0 | 0.0 | 0.0 | 0.43 | 0.65 | 0.62 | 0.73 | 0.73 |
| rp06 | 168 | 14.1 | 0.0 | 14.1 | 29.0 | 30.9 | 0.73 | 0.32 | 0.62 | 0.20 | 0.05 |
| rp07 | 185 | 24.6 | 24.9 | 24.9 | 26.1 | 24.9 | 0.52 | 0.16 | 0.50 | 0.09 | 0.11 |
| rp08 | 96 | 0.0 | 0.0 | 0.0 | 0.0 | 10.7 | 0.74 | 0.59 | 0.74 | 0.41 | 0.66 |
| rp10 | 171 | 0.0 | 10.4 | 10.3 | 10.3 | 9.3 | 0.88 | 0.73 | 0.71 | 0.47 | 0.63 |
| rp12 | 125 | 13.0 | 0.0 | 15.9 | 13.1 | 13.1 | 0.58 | 0.39 | 0.44 | 0.39 | 0.37 |
| rp13 | 71 | 0.0 | 0.0 | 0.0 | 14.8 | 8.4 | 0.84 | 0.01 | 0.84 | 0.50 | 0.59 |
| rp14_bound | 61 | 0.0 | 0.0 | 12.9 | 6.0 | 7.6 | 0.31 | 0.57 | 0.23 | 0.48 | 0.45 |
| rp14_free | 61 | 6.9 | 0.0 | 6.9 | 11.4 | 11.4 | 0.55 | 0.04 | 0.48 | 0.10 | 0.24 |
| rp15 | 68 | 0.0 | 0.0 | 15.5 | 19.8 | 16.3 | 0.52 | 0.27 | 0.57 | 0.41 | 0.45 |
| rp17 | 62 | 0.0 | 0.0 | 0.0 | 8.6 | 15.1 | 0.53 | 0.17 | 0.51 | 0.41 | 0.34 |
| rp19 | 71 | 0.0 | 0.0 | 0.0 | 16.3 | 12.5 | 0.58 | -0.15 | 0.54 | -0.01 | -0.01 |
| rp18 | 62 | 0.0 | 16.5 | 0.0 | 16.5 | 16.5 | 0.39 | 0.22 | 0.22 | -0.04 | -0.08 |
| rp21 | 41 | 0.0 | 0.0 | 0.0 | 18.0 | 18.0 | 0.02 | -0.28 | -0.06 | -0.41 | -0.51 |
| Average | | **3.5** | 4.0 | 6.0 | 12.1 | 12.4 | **0.54** | 0.30 | 0.48 | 0.27 | 0.30 |

Table S8. The RMSDs of structures with the lowest energy and Pearson correlation coefficients between energies and RMSDs of decoy structures calculated by the different statistical potentials for PM subset in test set III.

| | | RMSD of structure with the lowest energy | | | | | Pearson correlation coefficient (RMSD) | | | | |
|---|---|---|---|---|---|---|---|---|---|---|---|
| RNA | length | rsRNASP | RNA3DCNN | DFIRE-RNA | 3dRNAscore | RASP | rsRNASP | RNA3DCNN | DFIRE-RNA | 3dRNAscore | RASP |
| 1Z43 | 101 | 0.0 | 21.1 | 24.7 | 18.3 | 11.5 | 0.62 | 0.48 | 0.44 | -0.03 | 0.02 |
| 3A3A | 86 | 4.0 | 3.7 | 11.6 | 22.9 | 10.7 | 0.83 | 0.85 | 0.64 | 0.47 | 0.45 |
| 3IVN | 69 | 0.0 | 0.0 | 0.0 | 6.3 | 0.0 | 0.52 | 0.52 | 0.34 | 0.16 | -0.14 |
| 3L0U | 73 | 0.0 | 0.0 | 0.0 | 12.0 | 7.9 | 0.80 | 0.78 | 0.74 | -0.37 | -0.55 |
| 3LA5 | 71 | 0.0 | 0.0 | 0.0 | 0.0 | 20.8 | 0.87 | 0.59 | 0.85 | 0.04 | -0.33 |
| 3RKF | 67 | 0.0 | 0.0 | 1.0 | 9.5 | 11.2 | 0.96 | 0.71 | 0.97 | 0.48 | 0.49 |
| 3SKI | 68 | 0.0 | 0.0 | 0.0 | 9.9 | 11.0 | 0.89 | 0.83 | 0.82 | 0.18 | 0.20 |
| 4AOB | 94 | 0.0 | 0.0 | 0.0 | 11.6 | 18.6 | 0.73 | 0.75 | 0.63 | -0.50 | -0.34 |
| 4FEN | 67 | 0.0 | 0.0 | 0.0 | 8.8 | 22.2 | 0.89 | 0.64 | 0.89 | 0.05 | -0.27 |
| 5D5L | 77 | 0.0 | 0.0 | 0.0 | 19.6 | 11.4 | 0.66 | 0.44 | 0.48 | 0.29 | 0.26 |
| 5FJC | 93 | 0.0 | 0.0 | 0.0 | 20.3 | 0.0 | 0.84 | 0.74 | 0.83 | -0.38 | -0.06 |
| 5SWD | 65 | 0.0 | 0.0 | 0.0 | 5.1 | 6.0 | 0.78 | 0.29 | 0.38 | 0.39 | 0.36 |
| 6C27 | 47 | 8.0 | 10.0 | 9.1 | 8.0 | 9.3 | 0.55 | 0.64 | 0.45 | 0.59 | 0.65 |
| 6E8S | 38 | 0.0 | 12.3 | 11.3 | 12.2 | 11.3 | 0.29 | -0.06 | 0.34 | -0.10 | -0.07 |
| 6JQ6 | 81 | 23.3 | 28.6 | 22.6 | 23.2 | 22.6 | 0.12 | -0.30 | 0.00 | -0.33 | -0.34 |
| 6TFE | 52 | 0.0 | 0.0 | 9.6 | 0.0 | 11.8 | 0.37 | -0.04 | 0.38 | -0.05 | -0.29 |
| 6VMY | 148 | 0.0 | 0.0 | 33.4 | 19.7 | 21.8 | 0.36 | 0.35 | 0.26 | 0.35 | 0.33 |
| 7D82 | 50 | 14.6 | 12.5 | 7.1 | 12.5 | 14.6 | -0.10 | -0.09 | -0.12 | -0.39 | -0.38 |
| 7K16 | 51 | 0.0 | 0.0 | 0.0 | 4.7 | 4.7 | 0.73 | 0.10 | 0.71 | 0.43 | 0.41 |
| 7KJU | 75 | 0.0 | 0.0 | 13.8 | 13.1 | 16.9 | 0.72 | 0.68 | 0.65 | 0.50 | 0.58 |
| Average | | **2.5** | 4.4 | 7.2 | 11.9 | 12.2 | **0.62** | 0.44 | 0.53 | 0.09 | 0.05 |

Table S9. The DIs of structures with the lowest energy, ranks of the nearest-native structures (DIs) and Pearson correlation coefficients between energies and DIs of decoy structures calculated by the short- and long-range potentials of rsRNASP for Puzzles_standardized subset in test set III.

| | | DI of structure with the lowest energy | | | Rank of the nearest-native structure (DI) | | | Pearson correlation coefficient (DI) | | |
|---|---|---|---|---|---|---|---|---|---|---|
| RNA | length | Short-ranged | Long-ranged | Total | Short-ranged | Long-ranged | Total | Short-ranged | Long-ranged | Total |
| rp01 | 46 | 0.0 | 0.0 | 0.0 | 6 | 5 | 6 | 0.22 | 0.72 | 0.50 |
| rp02 | 100 | 2.8 | 0.0 | 0.0 | 6 | 2 | 4 | 0.66 | 0.44 | 0.65 |
| rp03 | 84 | 21.6 | 0.0 | 0.0 | 8 | 12 | 11 | 0.64 | 0.36 | 0.60 |
| rp04 | 126 | 5.0 | 4.7 | 5.0 | 15 | 21 | 16 | 0.44 | 0.53 | 0.49 |
| rp05 | 188 | 0.0 | 11.7 | 0.0 | 12 | 1 | 2 | 0.27 | 0.47 | 0.44 |
| rp06 | 168 | 19.2 | 20.5 | 19.2 | 3 | 4 | 2 | 0.66 | 0.74 | 0.78 |
| rp07 | 185 | 0.0 | 33.2 | 33.3 | 7 | 18 | 15 | 0.41 | 0.68 | 0.62 |
| rp08 | 96 | 0.0 | 0.0 | 0.0 | 3 | 17 | 13 | 0.57 | 0.79 | 0.70 |
| rp09 | 71 | 8.7 | 9.1 | 8.6 | 5 | 3 | 1 | 0.67 | 0.94 | 0.87 |
| rp10 | 171 | 0.0 | 0.0 | 0.0 | 2 | 12 | 7 | 0.93 | 0.91 | 0.94 |
| rp11 | 57 | 17.3 | 10.1 | 16.5 | 35 | 50 | 39 | -0.17 | -0.31 | -0.24 |
| rp12 | 125 | 18.1 | 15.9 | 17.8 | 37 | 29 | 38 | 0.62 | 0.71 | 0.83 |
| rp13 | 71 | 0.0 | 0.0 | 0.0 | 19 | 1 | 5 | 0.65 | 0.81 | 0.82 |
| rp14_bound | 61 | 0.0 | 14.5 | 0.0 | 15 | 33 | 19 | 0.49 | 0.14 | 0.43 |
| rp14_free | 61 | 0.0 | 8.1 | 0.0 | 1 | 4 | 1 | 0.34 | 0.55 | 0.66 |
| rp15 | 68 | 0.0 | 21.7 | 0.0 | 63 | 42 | 45 | 0.38 | 0.56 | 0.52 |
| rp17 | 62 | 11.6 | 16.7 | 0.0 | 58 | 3 | 22 | 0.35 | 0.62 | 0.54 |
| rp18 | 71 | 0.0 | 15.5 | 0.0 | 11 | 7 | 7 | 0.56 | 0.72 | 0.66 |
| rp19 | 62 | 18.6 | 0.0 | 0.0 | 26 | 7 | 13 | 0.35 | 0.08 | 0.23 |
| rp20 | 68 | 18.9 | 8.1 | 0.0 | 27 | 16 | 24 | 0.40 | 0.40 | 0.44 |
| rp21 | 41 | 34.2 | 8.2 | 0.0 | 23 | 3 | 4 | -0.14 | 0.70 | 0.48 |
| rp24 | 112 | 0.0 | 43.0 | 0.0 | 3 | 4 | 1 | 0.67 | 0.18 | 0.52 |
| Average | | 8.0 (11/22) | 11.0 (7/22) | **4.6 (16/22)** | 17.5 | **13.4** | **13.4** | 0.45 | 0.53 | **0.57** |

Table S10. The DIs of structures with the lowest energy, ranks of the nearest-native structure and Pearson correlation coefficients between energies and DIs of decoy structures calculated by the short- and long-ranges of rsRNASP for PM subset in test set III.

| | | DI of structure with the lowest energy | | | Rank of the nearest-native structure (DI) | | | Pearson correlation coefficient (DI) | | |
|---|---|---|---|---|---|---|---|---|---|---|
| RNA | length | Short-ranged | Long-ranged | Total | Short-ranged | Long-ranged | Total | Short-ranged | Long-ranged | Total |
| 1Z43 | 101 | 0.0 | 14.7 | 0.0 | 17 | 28 | 22 | 0.56 | 0.51 | 0.66 |
| 3A3A | 86 | 4.8 | 32.6 | 4.8 | 3 | 26 | 6 | 0.88 | -0.12 | 0.85 |
| 3IVN | 69 | 0.0 | 0.0 | 0.0 | 2 | 6 | 1 | 0.57 | 0.36 | 0.48 |
| 3L0U | 73 | 0.0 | 0.0 | 0.0 | 2 | 11 | 1 | 0.60 | 0.83 | 0.80 |
| 3LA5 | 71 | 0.0 | 0.0 | 0.0 | 2 | 1 | 1 | 0.48 | 0.92 | 0.88 |
| 3RKF | 67 | 0.0 | 1.1 | 0.0 | 1 | 12 | 3 | 0.86 | 0.95 | 0.95 |
| 3SKI | 68 | 0.0 | 0.0 | 0.0 | 4 | 4 | 2 | 0.81 | 0.76 | 0.89 |
| 4AOB | 94 | 0.0 | 0.0 | 0.0 | 2 | 1 | 1 | 0.66 | 0.74 | 0.78 |
| 4FEN | 67 | 0.0 | 0.0 | 0.0 | 4 | 4 | 2 | 0.71 | 0.86 | 0.86 |
| 5D5L | 77 | 0.0 | 0.0 | 0.0 | 8 | 14 | 10 | 0.76 | 0.48 | 0.72 |
| 5FJC | 93 | 0.0 | 0.0 | 0.0 | 1 | 1 | 1 | 0.72 | 0.87 | 0.86 |
| 5SWD | 65 | 6.7 | 44.8 | 0.0 | 4 | 8 | 3 | 0.76 | -0.13 | 0.69 |
| 6C27 | 47 | 11.4 | 11.7 | 9.7 | 8 | 10 | 5 | 0.68 | 0.11 | 0.60 |
| 6E8S | 38 | 21.1 | 0.0 | 0.0 | 34 | 23 | 30 | -0.03 | 0.44 | 0.26 |
| 6JQ6 | 81 | 0.0 | 31.5 | 33.3 | 4 | 16 | 7 | 0.27 | 0.00 | 0.21 |
| 6TFE | 52 | 8.3 | 0.0 | 0.0 | 32 | 27 | 35 | 0.18 | 0.44 | 0.38 |
| 6VMY | 148 | 0.0 | 57.7 | 0.0 | 7 | 35 | 13 | 0.74 | -0.56 | 0.47 |
| 7D82 | 50 | 17.7 | 7.0 | 17.7 | 5 | 23 | 10 | -0.07 | 0.20 | -0.03 |
| 7K16 | 51 | 0.0 | 0.0 | 0.0 | 3 | 3 | 2 | 0.77 | 0.38 | 0.73 |
| 7KJU | 75 | 19.8 | 52.9 | 0.0 | 13 | 19 | 7 | 0.80 | -0.09 | 0.66 |
| Average | | 4.5 (13/20) | 12.7 (11/20) | **3.3 (16/20)** | **7.8** | 13.6 | 8.1 | 0.59 | 0.40 | **0.64** |

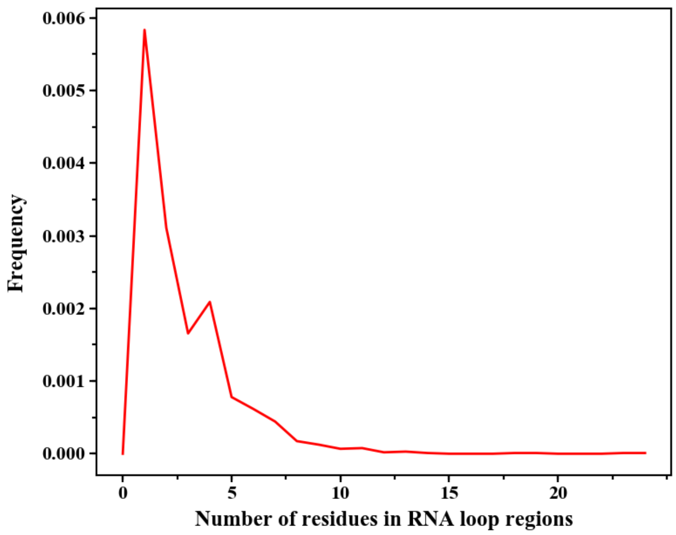


Figure S1. Frequency of the number of residues in RNA loop regions for 191 native RNA structures in the training set; see the details in Section 2 in the Supplementary Material. Here, the secondary structures for 191 native RNA structures were derived with x3DNA-DSSR (5).

15

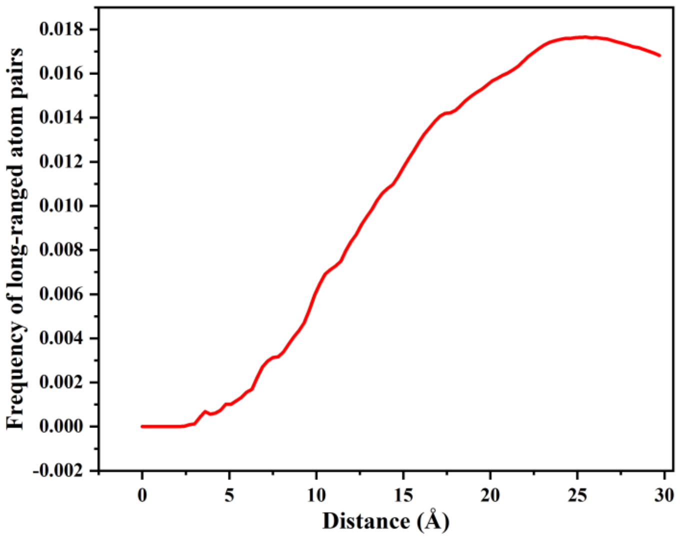


Figure S2. Frequency of long-ranged atom pairs for 191 native RNAs in the training set; see the details in Section 3 in the Supplementary Material.

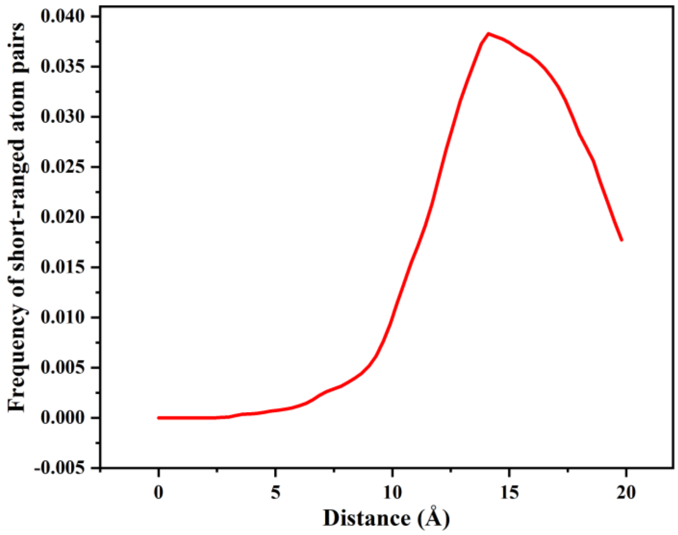

Figure S3. Frequency of short-ranged atom pairs for 191 native RNAs in the training set; see the details in Section 3 in the Supplementary Material.

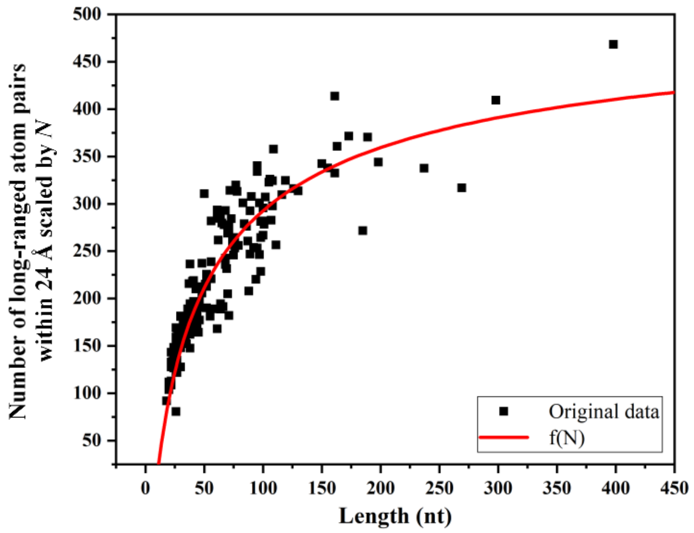


Figure S4. Relationship between number of long-ranged atom pairs within 24 Å scaled by $N$ (per nucleotide) and RNA length ($N$, in nt) for 191 native RNAs in the native training set; see the details in Section 4 in the Supplementary Material. Here, $f(N) = \frac{-2685}{(N+16)^{0.5}} + 542$ denotes the fitted line.

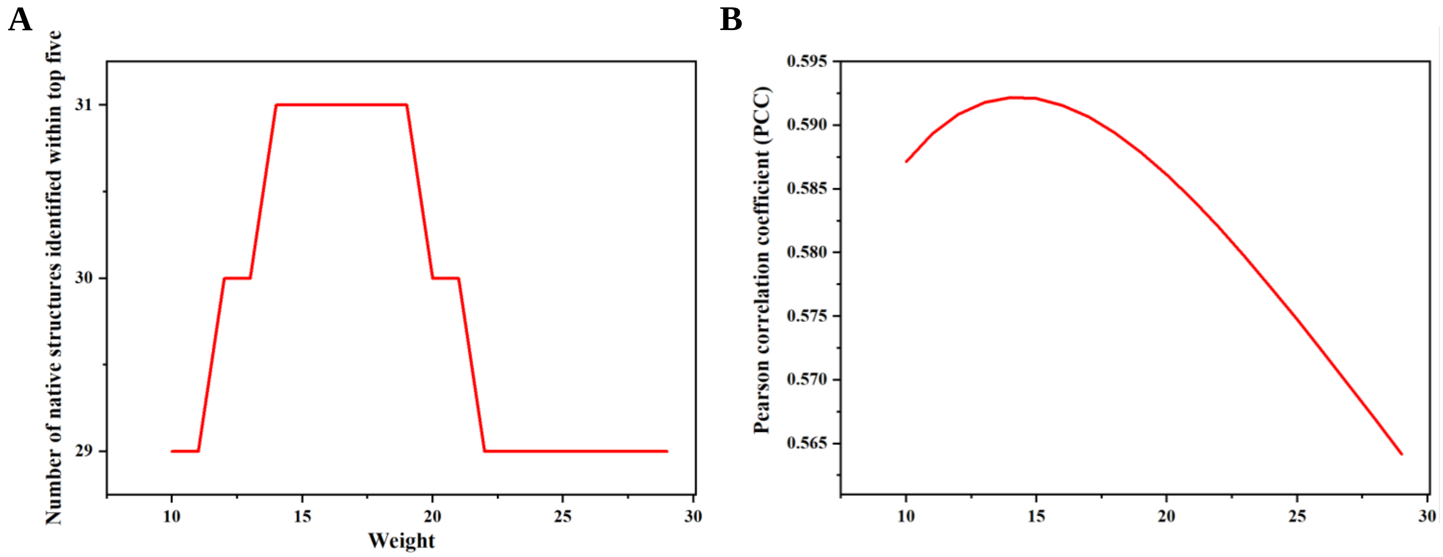


Figure S5. (A) Number of native structures identified within top five of the lowest energies for the training decoy set under different weight $w_0$; (B) Pearson correlation coefficient for the training decoy set under different weight $w_0$. Here, the training decoy set is composed of 35 single-stranded RNAs with about 40 decoy structures for each RNA from the four 3D structure prediction models (6-9); see the details in Section 4 in the Supplementary Material.

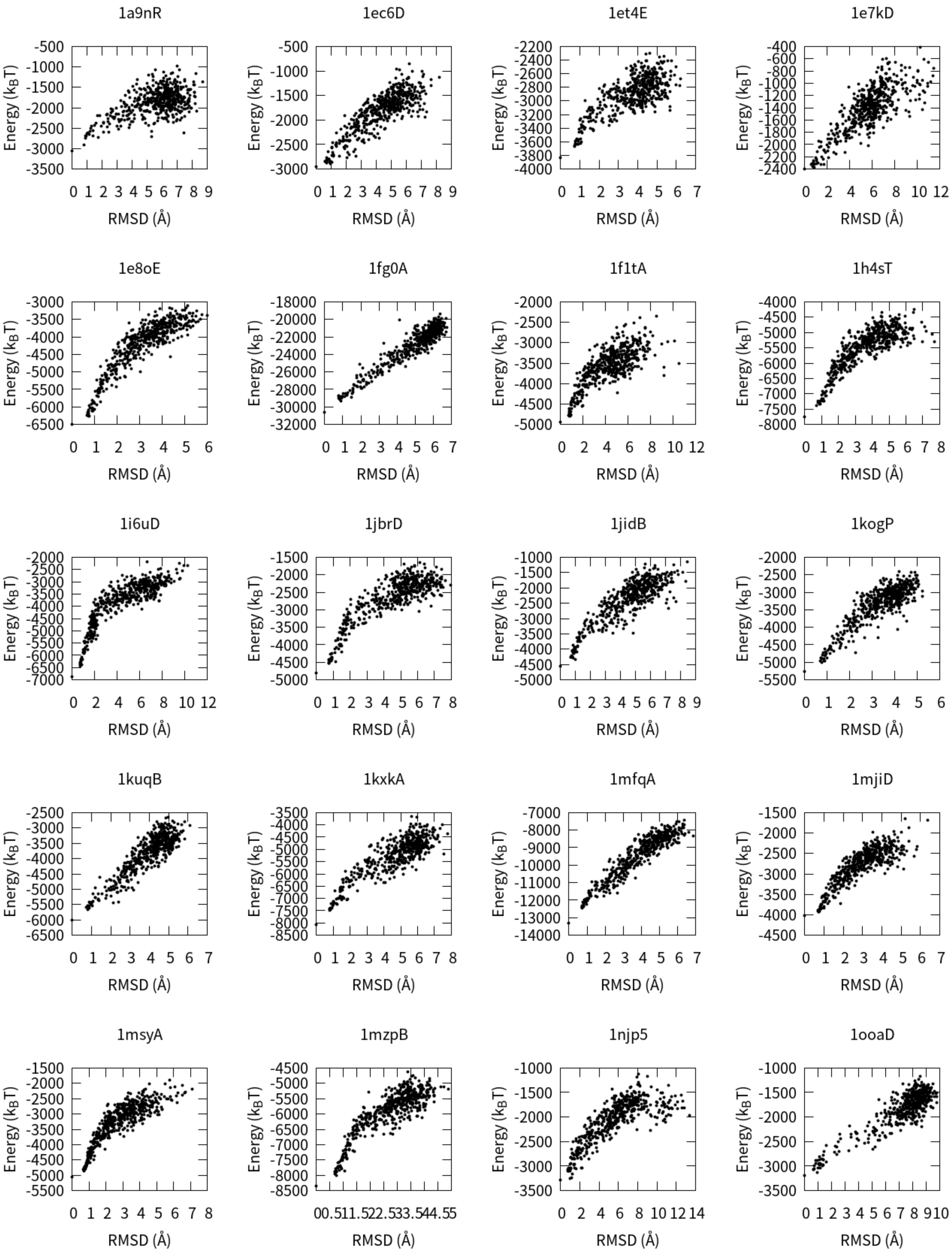
1a9nR
1ec6D
1et4E
1e7kD
1e8oE
1fg0A
1f1tA
1h4sT
1i6uD
1jbrD
1jidB
1kogP
1kuqB
1kxkA
1mfqA
1mjiD
1msyA
1mzpB
1njp5
1ooaD
Energy ($k_BT$)
RMSD (Å)

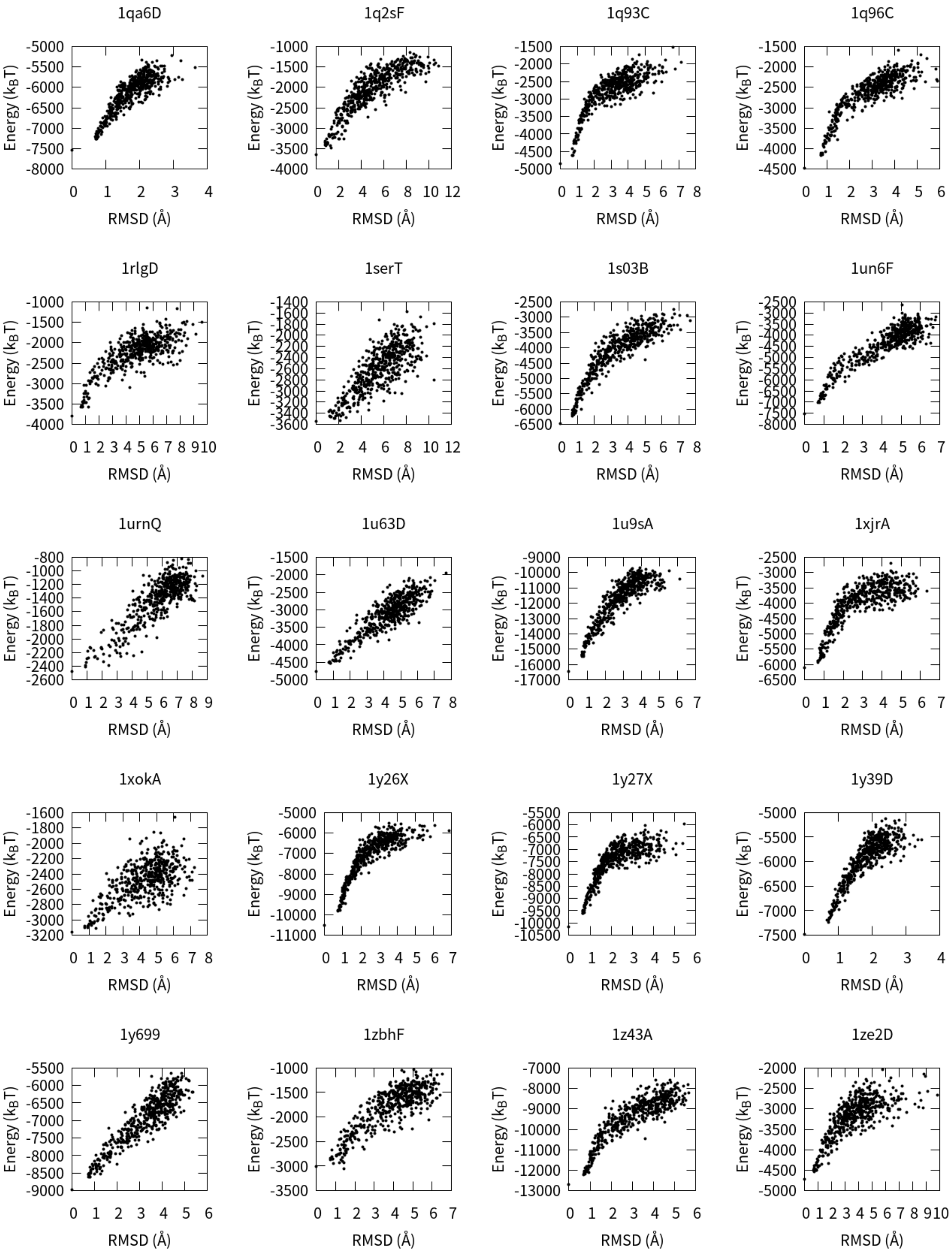
1qa6D
1q2sF
1q93C
1q96C
1rlgD
1serT
1s03B
1un6F
1urnQ
1u63D
1u9sA
1xjrA
1xokA
1y26X
1y27X
1y39D
1y699
1zbhF
1z43A
1ze2D
Energy ($k_BT$)
RMSD (Å)

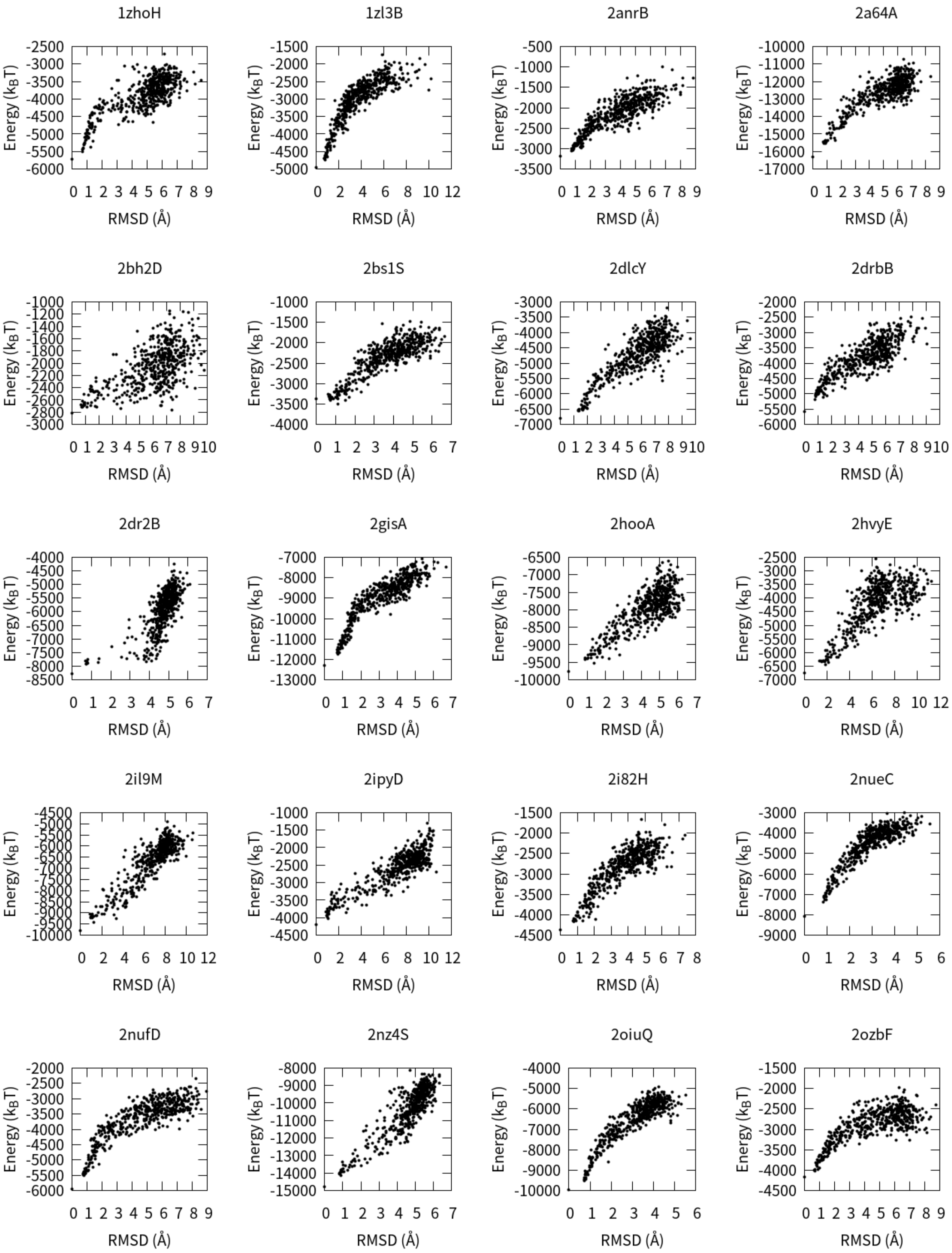
1zhoH
1zl3B
2anrB
2a64A
2bh2D
2bs1S
2dlcY
2drbB
2dr2B
2gisA
2hooA
2hvyE
2il9M
2ipyD
2i82H
2nueC
2nufD
2nz4S
2oiuQ
2ozbF
Energy (kBT)
RMSD (Å)

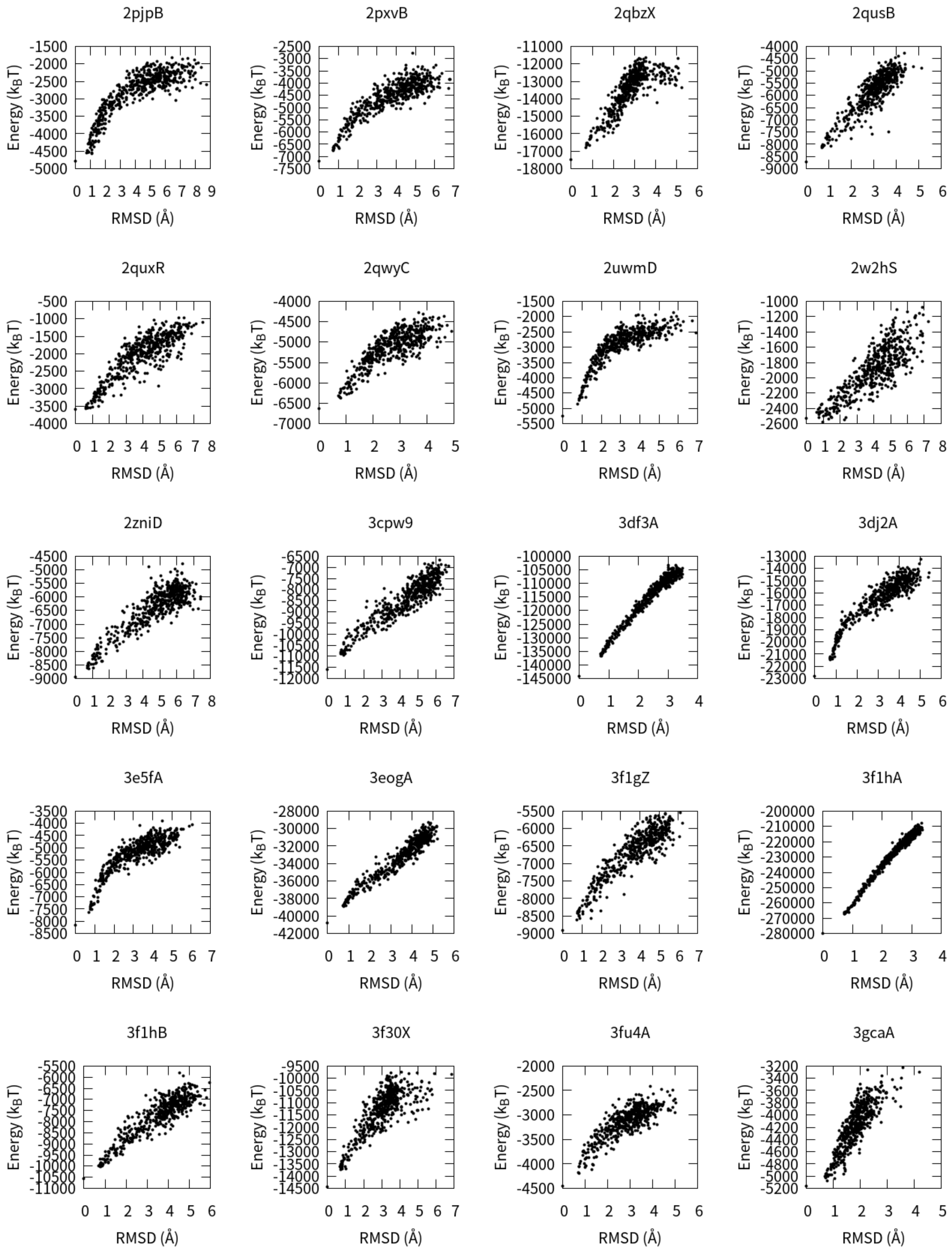
2pjpB
2pxvB
2qbzX
2qusB
2quxR
2qwyC
2uwmD
2w2hS
2zniD
3cpw9
3df3A
3dj2A
3e5fA
3eogA
3f1gZ
3f1hA
3f1hB
3f30X
3fu4A
3gcaA
Energy ($k_BT$)
RMSD (Å)

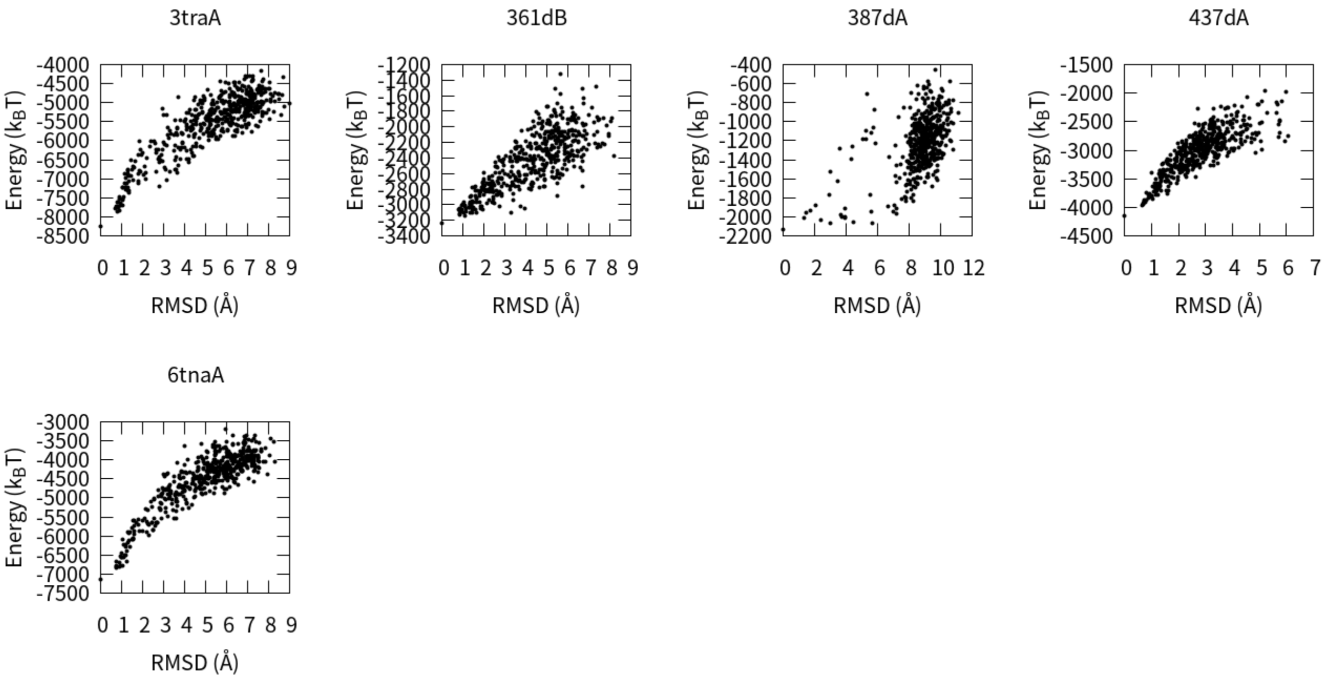


Figure S6. The RMSD-energy scatter-plots for all the 85 RNAs in test set I by rsRNASP.

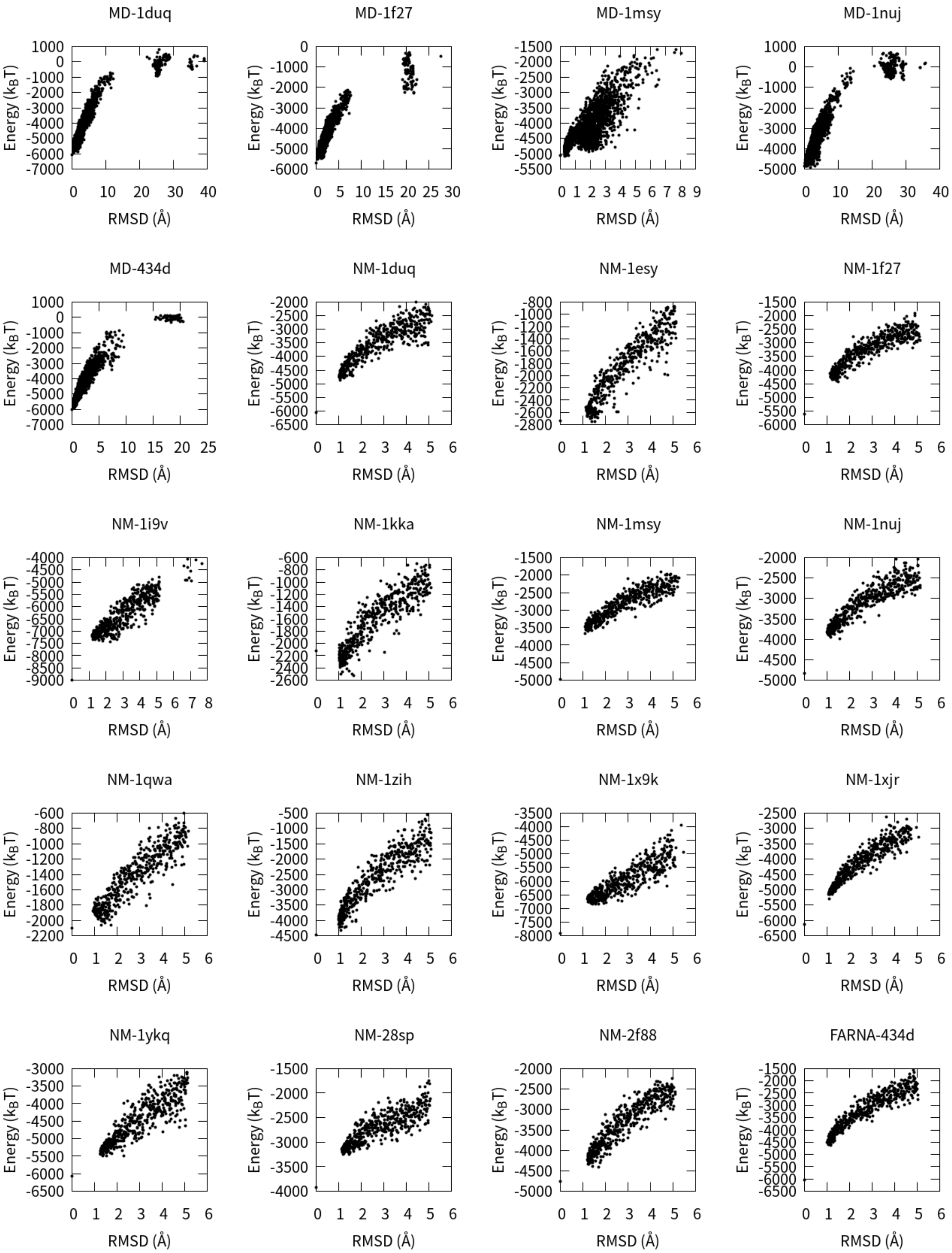
MD-1duq
MD-1f27
MD-1msy
MD-1nuj
MD-434d
NM-1duq
NM-1esy
NM-1f27
NM-1i9v
NM-1kka
NM-1msy
NM-1nuj
NM-1qwa
NM-1zih
NM-1x9k
NM-1xjr
NM-1ykq
NM-28sp
NM-2f88
FARNA-434d
Energy ($k_BT$)
RMSD (Å)

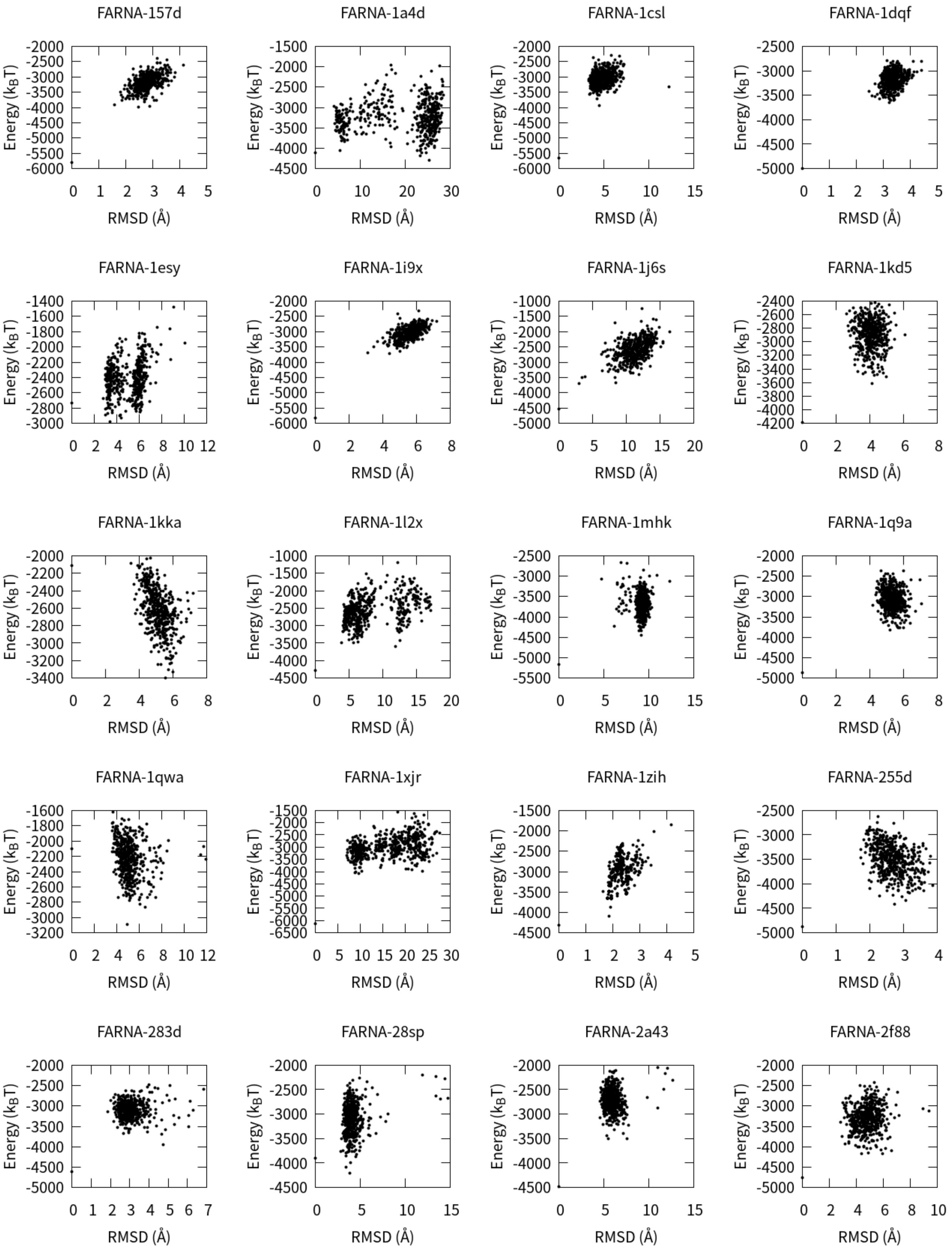


Figure S7. The RMSD-energy scatter-plots for all the 40 RNAs in test set II by rsRNASP.

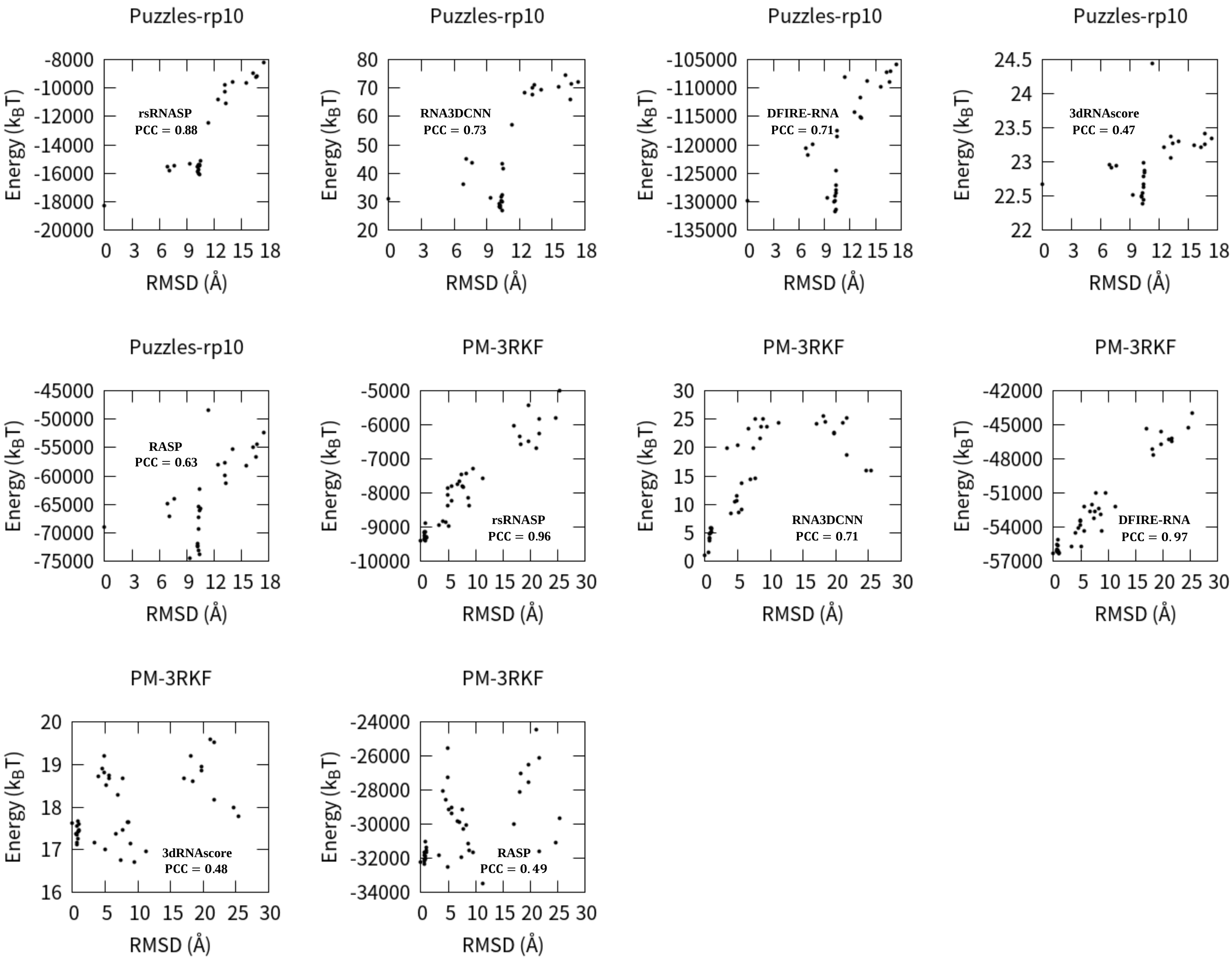


Figure S8. The RMSD-energy scatter-plots for four examples (puzzle-10 in Puzzles_standardized subset; 3RKF in PM subset) given by five scoring functions (rsRNASP, RNA3DCNN, DFIRE-RNA, 3dRNAscore and RASP). The PCC values on RMSDs are also annotated in each panel.

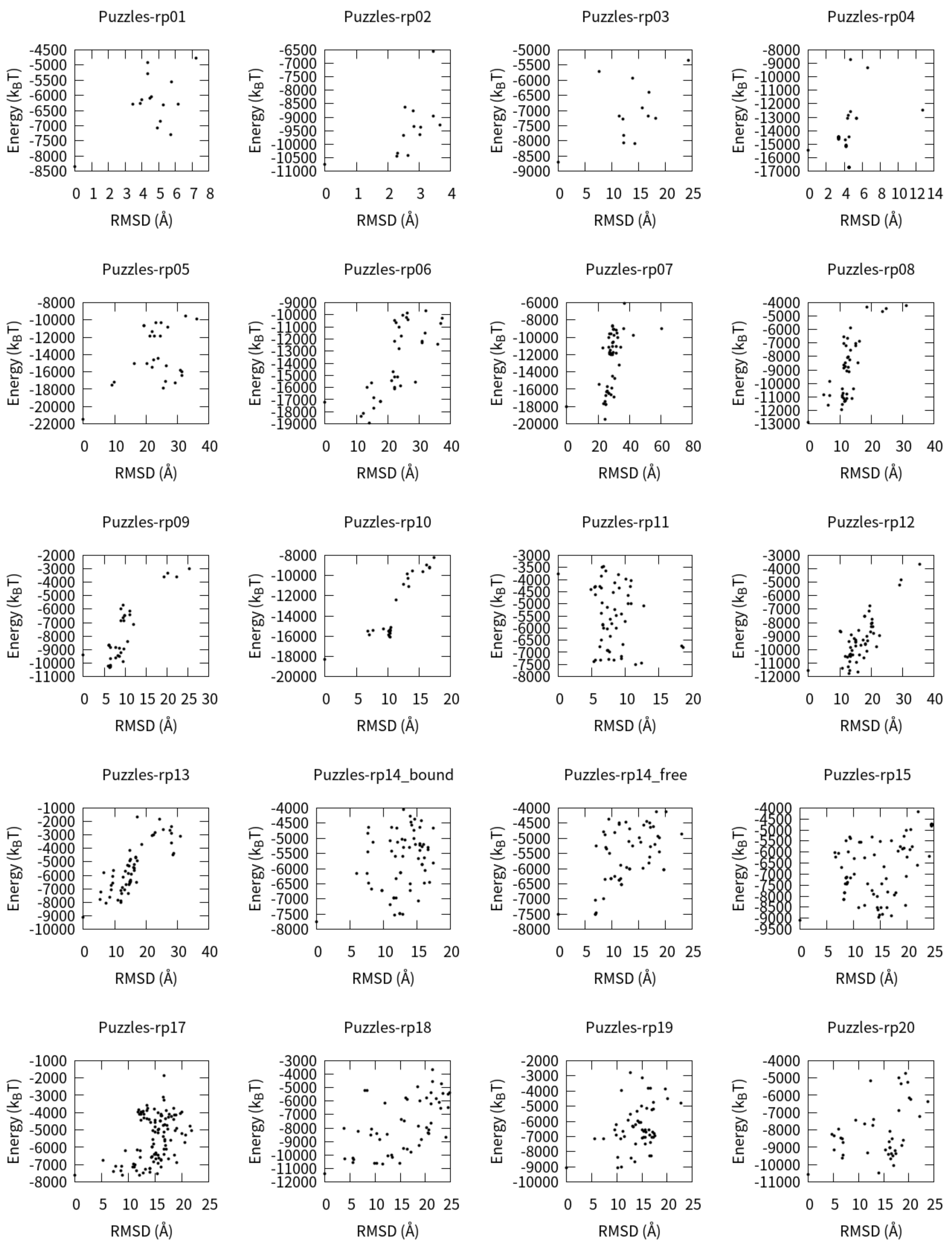
Puzzles-rp01
Puzzles-rp02
Puzzles-rp03
Puzzles-rp04
Puzzles-rp05
Puzzles-rp06
Puzzles-rp07
Puzzles-rp08
Puzzles-rp09
Puzzles-rp10
Puzzles-rp11
Puzzles-rp12
Puzzles-rp13
Puzzles-rp14_bound
Puzzles-rp14_free
Puzzles-rp15
Puzzles-rp17
Puzzles-rp18
Puzzles-rp19
Puzzles-rp20
Energy ($k_BT$)
RMSD (Å)

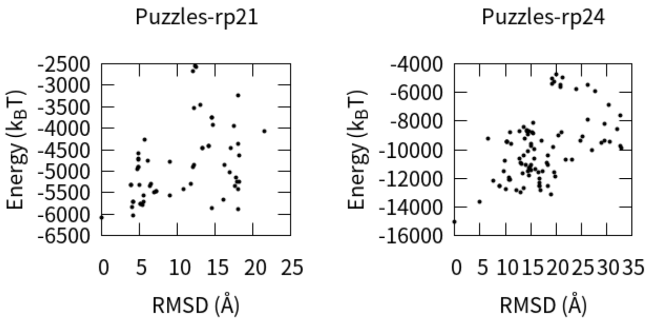


Figure S9. The RMSD-energy scatter-plots for all 22 puzzles in test set III_ Puzzles_standardized subset by rsRNASP.

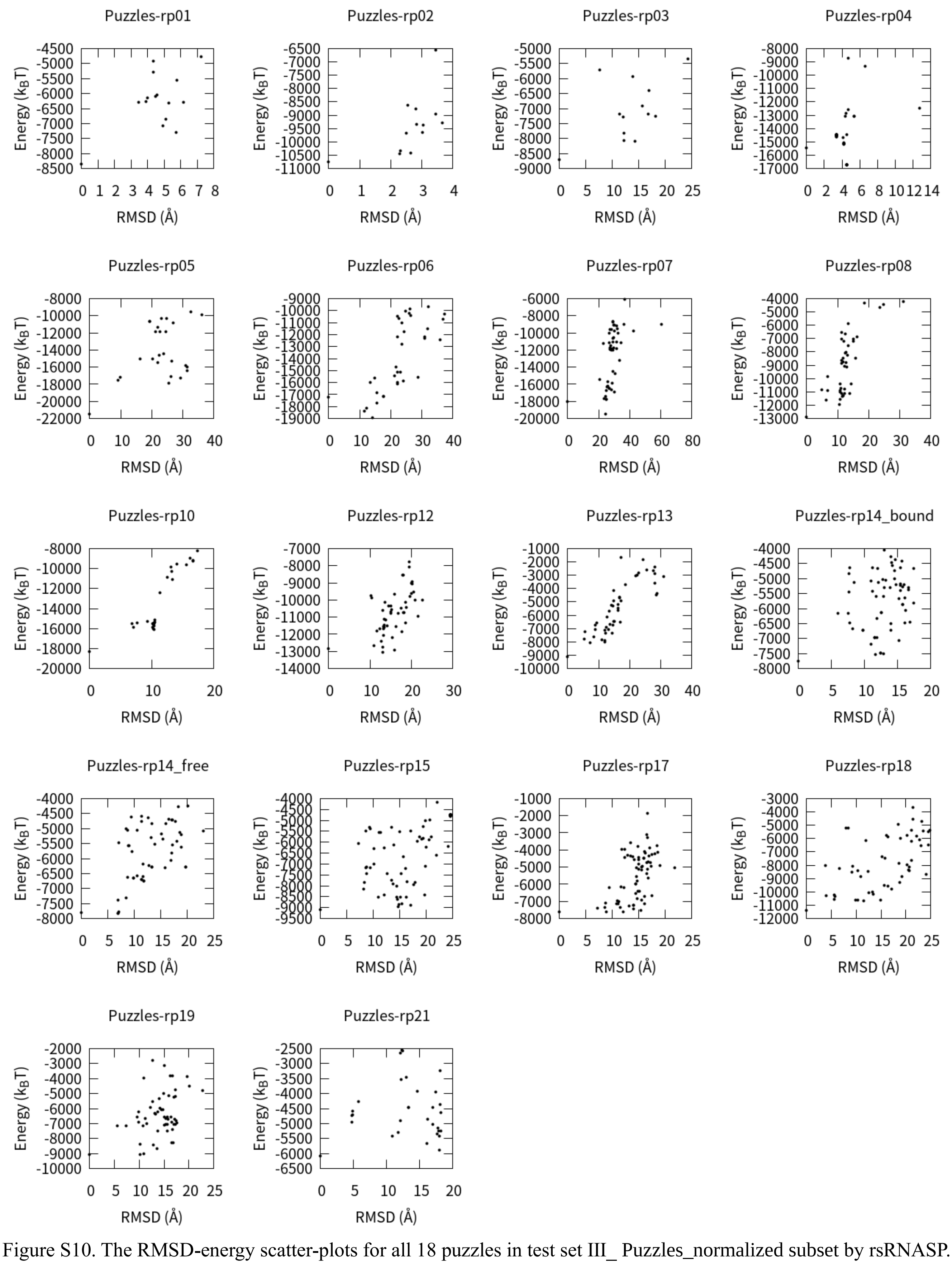

Figure S10. The RMSD-energy scatter-plots for all 18 puzzles in test set III_ Puzzles_normalized subset by rsRNASP.

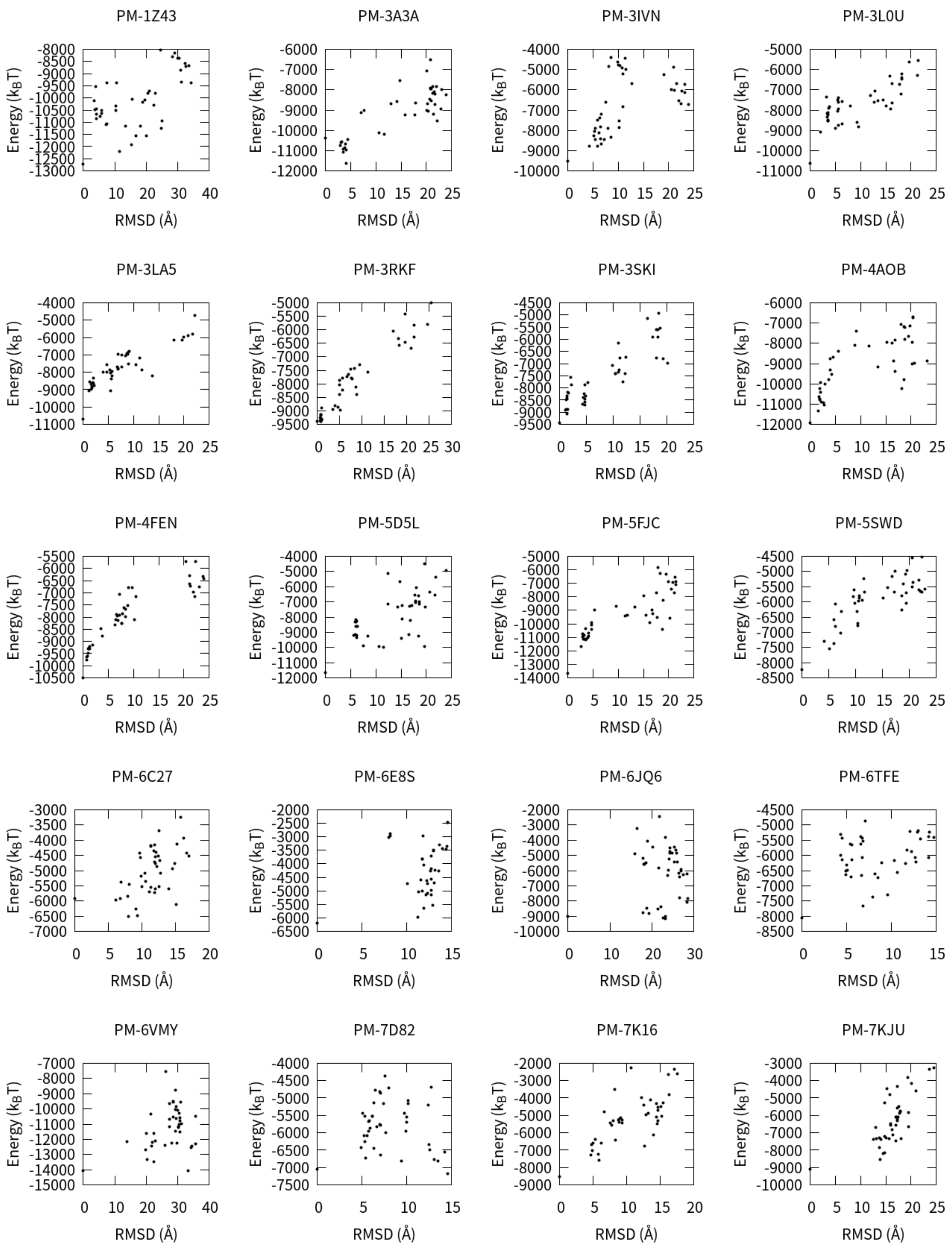


Figure S11. The RMSD-energy scatter-plots for all 20 puzzles in test set III_PM subset by rsRNASP.

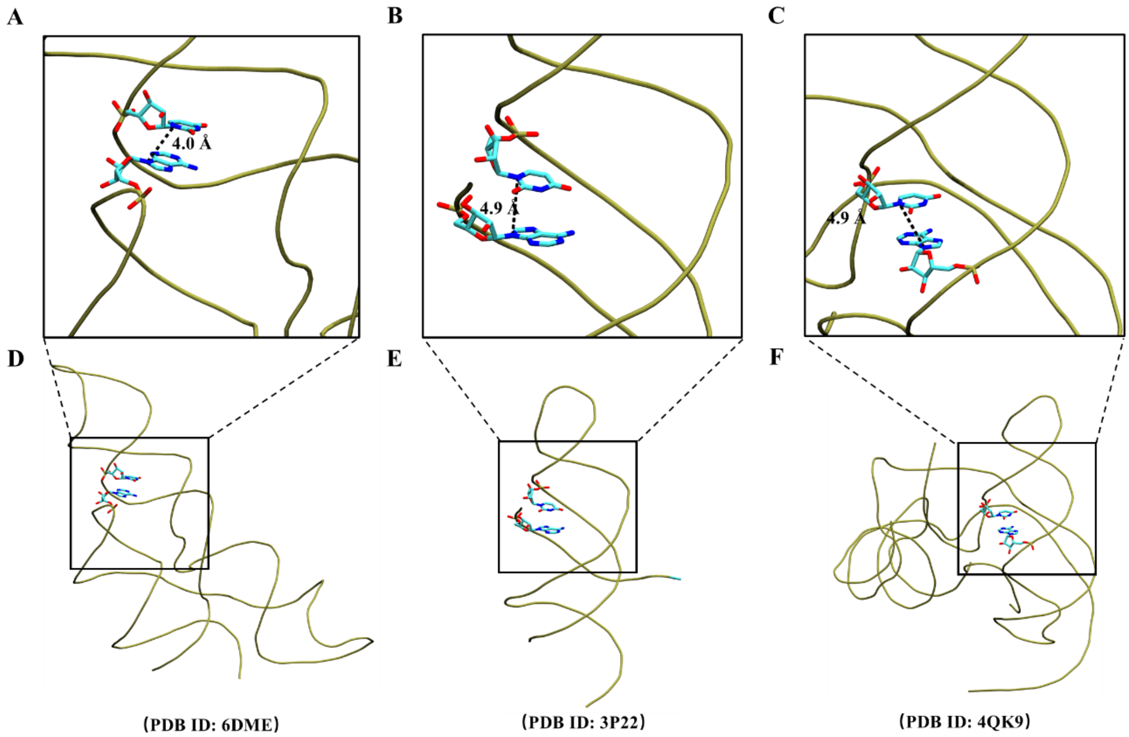


Figure S12. (A)-(C) Representative distances between AN9 and UN1 for base-stacking interaction of two residues in adjacent branches of RNA 3D structure (PDB ID: 6DME), base-stacking interaction of triple-helical region (PDB ID: 3P22), and coaxial-stacking interaction at junction region (PDB ID: 4QK9) in the long-ranged potential, respectively. The bottom panels show the corresponding complete 3D structures for the local structures in panels (D)-(F).
